# Ependymocytes control cerebrospinal fluid flow to the peripheral organs through periaxonal pathway

**DOI:** 10.1101/2023.10.09.560634

**Authors:** Xinyu Li, Siman Wang, Dianjun Zhang, Yuliang Feng, Yingyu Liu, Weiyang Yu, Shu Li, Lulu Cui, Tibor Harkany, Alexei Verkhratsky, Maosheng Xia, Baoman Li

## Abstract

Mechanisms controlling movement of the CSF through the central canal towards the peripheral nerves are poorly characterized. We found that fluorescent dyes injected into cisterna magna are carried with cerebrospinal fluid (CSF) through the central canal and peripheral nerves to the peripheral organs such as liver, and pancreas. We also found close connection between spinal axons and ependymocytes, suggesting synaptic interactions. Serotonin, acting through the 5-HT_2B_ receptors abundantly expressed in ependymal cells, trigger Ca^2+^ signal that induces polymerization of cytoskeleton protein F-actin, consequently reducing the volume of ependymocytes. Shrinkage of the latter opens one-way route to facilitate CSF outflow from the central canal into the spinal cord parenchyma and peripheral nerves. In liver, CSF is received by stellate cells. Ependymal control over transfer of the CSF from central canal to peripheral organs by the periaxonal space (PAS) represents a novel mechanism dynamically connecting the CNS with the periphery.

**In brief:** Ependymocytes control CSF flow from CNS to peripheral organs by periaxonal pathway, and serotonin evokes the ependymal shrinkage by the aggregation of F-actin.

**Highlights:** Ependymocytes control CSF flow from central canal to peripheral organs; CSF flows through the peripheral periaxonal space to reach the peripheral organs; Serotonin makes ependymocytes shrunk by 5-HT_2B_ receptor-mediated Ca^2+^ signaling and F-actin polymerization; The hepatic stellate cells are potential collectors of CSF in the liver.

## Introduction

The cerebrospinal fluid (CSF) is a weak salt solution that fills in and circulates through the ventricles, the central canal, the subarachnoid space and reaches peripheral organs along cranial and spinal nerves. The CSF is mainly produced by choroid plexi, located in the lateral ventricles and third and fourth ventricles. Average CSF volume in mice is around 50 μl, whereas in humans CSF volume ranges between 150 and 300 ml ^1–4^. The CSF in humans is produced at a rate of ∼500 - 600 ml per day (or ∼ 350 μl per minute), thus being fully replaced approximately every 6 hours ^5^. The CSF has many functions ^5,6^, in particular, the CSF provides a pathway for long-range volume transmission mediated by metabolites and neuroactive substances including neurotransmitters, peptides, extracellular vesicles, and immunocompetent molecules ^7^. Neurotransmitters and neurohormones are secreted to the CSF to mediate long-range signaling ^8^. Signaling through the CSF may even reach peripheral organs through cranial and peripheral nerves that form the peripheral CSF fluid outflow pathway ^9,10^, which, howvere, was not studied in detail hitherto. Similarly unknown are the mechanisms regulating the CSF flow from the CNS structures to the peripheral nerves and hence to peripheral organs.

Here we identified ependymocytes to dynamically modulate intercellular conduit for CSF drainage from the spinal central canal to the peripheral organs. Ependymal glia are direct scions of radial glia, being thus a part of astroglia, a class of homeostatic cells of the brain ^11^. Ependymal glia express astroglial marker vimentin, and are connected into a functional syncytium through gap junctions ^12,13^. Ependymocytes display many physiological properties of astroglia, being electrically non-excitable and relying on intracellular ionic (mainly Ca^2+^) signalling, that propagates through ependymal syncytium. In addition, ependymal cells are endowed with neurotransmitter receptors, while many of ependymocytes receive synaptic contacts from neurones ^8^. We found that stimulation of ependymal cells of the spinal canal with serotonin triggers rapid contraction of cytoskeletal F-actin leading to a volumetric decrease of ependymocytes, which opens the route for CSF flow from the CNS to peripheral nerves and peripheral organs.

## Results

### Identification of CSF flow from ventricles to liver

A visible blue dye cadaverine (1 kDa) was injected into cisterna magna (CIM) at 5 μl (20 mg/ml), as shown in Fig. 1A. The injected cadaverine distributed from CIM to the brain, spinal cord, nerve branches, and subsequently appeared in liver and pancreas (Fig. 1B). Blue cadaverine staining propagated from the CIM through central canal to cauda equina in 30 minutes (Fig. 1B) and could be detected in liver and pancreas 4 hours after intra-CIM injection (Fig. S1A). After 12 hours, cadaverine staining disappears from both CNS and peripheral organs (Fig. 1B). To further characterize CSF flow from CNS to the periphery, the fluorescence dyes FITC-D40 (40 kDa) and TR-D3 (3 kDa) were co-injected into CIM. Four hours after injection, the distribution of FITC-D40 and TR-D3 fluorescence was monitored in the brain (anterior section: bregma 0.62 mm and posterior section: −1.34 mm), in the brain stem (Bregma −6.12 mm), in the cervical (C3), thoracic (T5) and lumbar (L3) sections of the spinal cord and in the liver and pancreas, as shown in Fig. 1C-E and Fig. S1. In the liver 4 hours after the injection, the red fluorescence tracer TR-D3 was localized in the hepatic stellate cells; the latter were labeled either with vimentin (Fig. 1D) or with α-smooth muscle actin (α-SMA, Fig. 1E), which is the cytoskeletal protein ^14^ used as a marker of hepatic stellate cells activation ^15^. Similarly, the fluorescence tracer TR-D3 was detected in pancreatic stellate cells stained with α-SMA (Fig. S1B).

**Figure 1.**
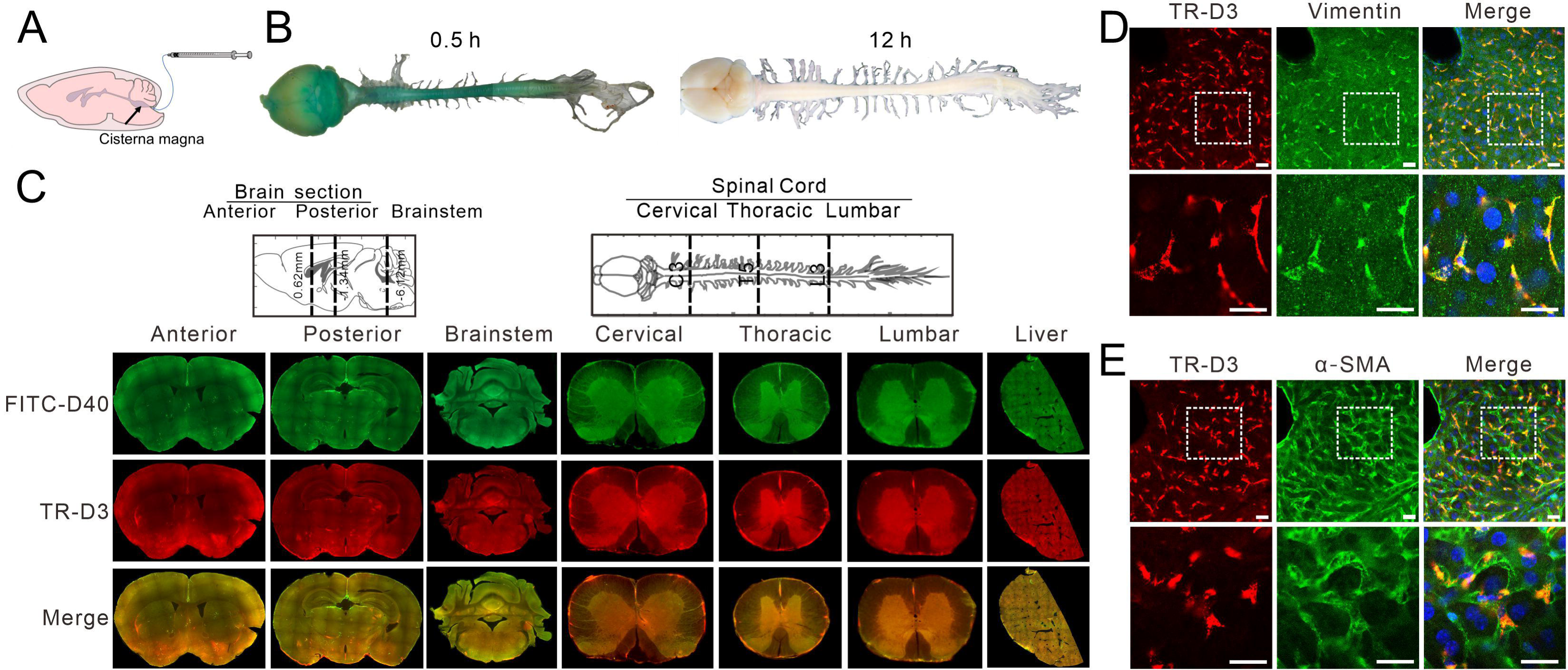
Delivery of CSF from cisterna magna into liver. A: The schematic of tracer injection into cisterna magna (CIM) at 5 μl (20 mg/ml). B: Left: Images of the brain and spinal cord 30 min after cadaverine CIM-injection. Right panel: same image 12 hours after injection. C: Representative images of fluorescence tracers FITC-D40 (green) and TR-D3 (red) in anterior and posterior brain sections, brainstem, cervical, thoracic and lumbar spinal cord, and in the liver. D: Representative images of TR-D3 fluorescence (red) in liver co-stained with vimentin (green) and nucleus marker DPAI (blue). Scale bar = 20 μm. E: Representative images of fluorescence TR-D3 (red) in liver co-stained with α-SMA (green) and nucleus marker DPAI (blue). Scale bar = 20 μm.

To exclude CSF refluxing by superior vena cava by the way of meningeal lymphatic vessels ^16^ or through anastomoses between CSF and vessels by fenestrations in perivascular space ^17^, and confirm the primary contribution of the spinal nerve route, the head branch of the superior vena cava was ligated or the spinal nerves were sectioned, respectively, and the emergence of the tracer in the liver was monitored (Fig. 2A-C). When the head branch of the superior vena cava, which collects the lymph and blood from the brain, was ligated (Fig. S2A), the level of CIM injected TR-D3 fluorescence in the liver decreased to 73.87% ± 5.97% of the control group (p = 0.012; n = 6; Fig. 2C). Conversely, sectioning of the spinal nerves (T2-T12) innervating the liver (Fig. S2B), decreased the fluorescence intensity of TR-D3 delivered after injection to CIM to 30.93% ± 6.42% of the control group (p < 0.001; n = 6; Fig. 2C). To further characterize the flow routes of CSF along spinal nerves, Thy1-YFP transgenic mice in which neuronal soma and axons are randomly fluorescently labeled with YFP were CIM-injected with TR-D3 in anaesthetized mice. The spinal nerves (T7-T9) were surgically exposed and imaged with two-photon microscopy. The tracer reached the nerves in about 35 – 55 minutes after injection and its movement along the nerve was captured (Fig. 2D and 2E; supplementary video 1).

**Figure 2.**
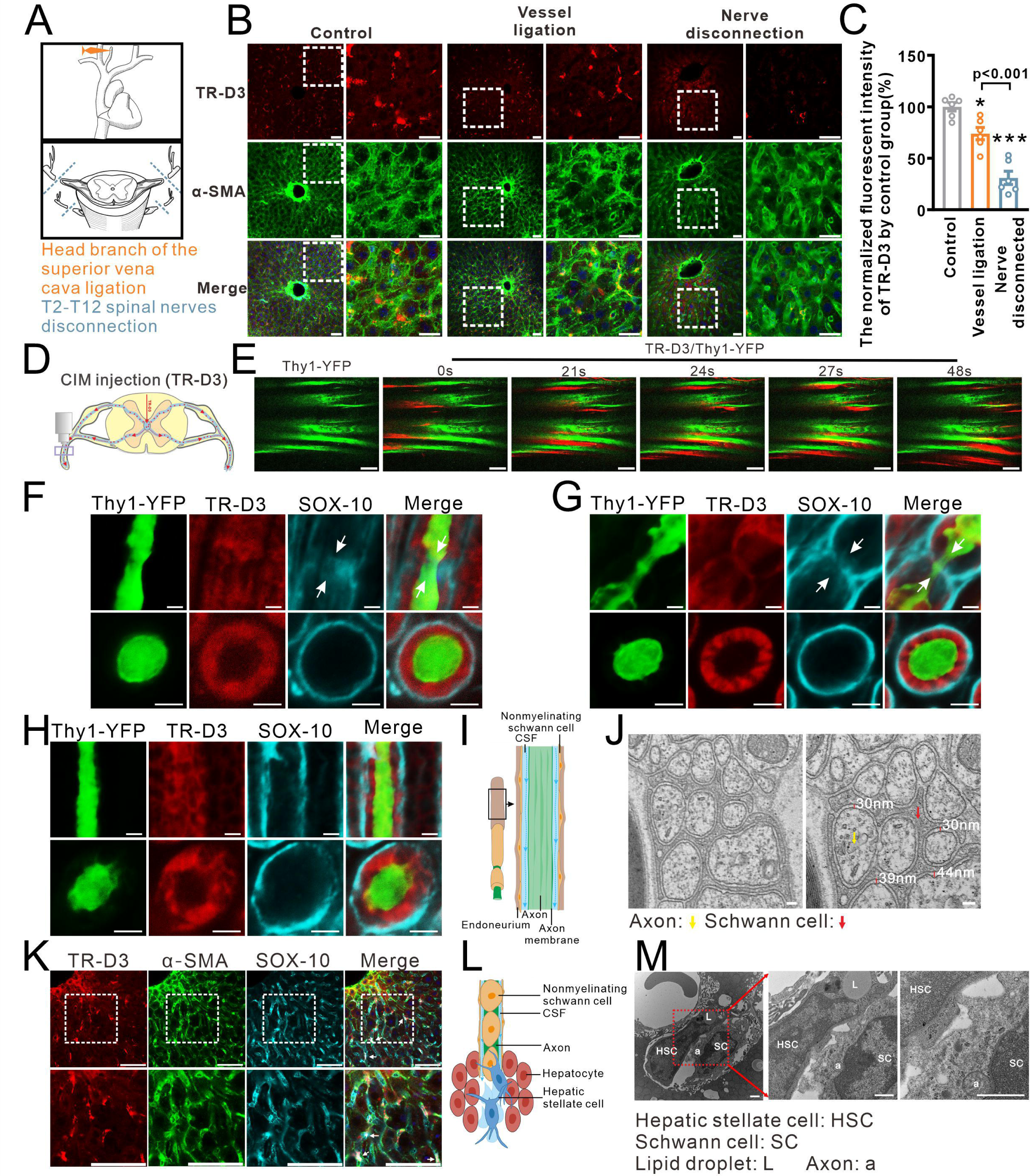
CSF connects to periaxonal pathway. A: Upper panel: the schematic of the head branch of superior vena cava ligation. Below panel: the schematic of the spinal T2-T12 nerves disconnection. B: After the operation of superior vena cava ligation or spinal nerves disconnection, TR-D3 was injected into CIM for 4 hours, the liver tissues were observed. Representative fluorescence images of TR-D3 (red) distributed in liver co-stained with α-SMA (green) after ligation of the head branch of superior vena cava or cutting off the spinal nerves T2-T12 segments. Scale bar=25 μm. C: The fluorescent intensity of TR-D3 normalized by control group. Open circles represent data from individual mice (n=6) and bars represent averages±SEM, as comparison with control group *p<0.05, ***p<0.001. D: Anatomical location of observation window for imaging spinal nerves. E: The images of red TR-D3 distributing along green YFP+ labeled axon of the spinal nerves in Thy1-YFP transgenic mice. Scale bar=100 μm. F: Representative images of longitudinal and cross sections of TR-D3 fluorescence (red) and sox-10 (green) of spinal nerves. Scale bar=5 μm. G: Representative images of longitudinal and cross sections of TR-D3 fluorescence (red) and sox-10 (green) of sciatic nerve. Scale bar= 5 μm. H: Representative longitudinal and cross section images of fluorescence tracer TR-D3 (red), sox-10 (cyan) and Thy1-YFP (green) in the nerve roots separated from the hepatic nerve plexus, 4 hours after TR-D3 was injected into CIM. Scale bar= 5 μm. I: The schematic diagram of CSF flowing in the periaxonal space. J: The representative electron microscope images of axon (yellow arrow) and Schwann cell (red arrow) in non-myelin axon of the hepatic nerve plexus. The red bar was the measured distance. Scale bar=100 nm. K: The representative images of fluorescence tracer TR-D3 (red) co-stained with α-SMA (green), sox-10 (cyan) and DAPI (blue) in the liver 4 hours after TR-D3 was injected into CIM at. The overlapping images of TR-D3, α-SMA and sox-10 are indicated by arrows. Scale bar=100 μm. L: The schematic diagram of the proposed connection between Schwann cells and hepatic stellate cells. M: The representative electron microscope images of hepatic stellate cell (HSC), Schwann cell (SC), axon (a) and lipid droplet (L) in liver. Scale bar=1 μm.

We further used Thy1-YFP transgenic mice to identify possible compartment(s) for CSF flow in the nerves, again employing TR-D3 as the fluorescence tracer. Four hours after intra-CIM injection of TR-D3 the spinal nerves (T7-T9; Fig. 2F) and the sciatic nerves (Fig. 2G) were co-stained with Schwann cell marker sox-10. In the longitudinal section (upper row) and the cross section (lower row), TR-D3 fluorescence appeared in close association with YFP green fluorescent axons and Schwann cells, both in spinal (Fig. 2F) and sciatic nerves (Fig. 2G). Thus, TR-D3 is flowing through periaxonal space limited with endoneurium. Moreover, the tracers were even delineating nodes of Ranvier (Fig. 2F and 2G), indicating that the tracer did not move outside of the periaxonal space delineated with endoneurium.

Liver is innervated mainly by non-myelinating axons of sympathetic splanchnic and parasympathetic vagal nerves, as well as by branches of right phrenic nerves containing both sympathetic and somatic spinal axons from T7-T9. Furthermore, the red TR-D3 was in close association with YFP green fluorescent axons and the non-myelinating sox-10 positive Schwann cells (Fig. 2H) indicating CSF flow along the periaxonal space (PAS) as shown in the schematic diagram on Fig. 2I. The electron microscope (EM) images of non-myeline also supported the existence of PAS, the size of the gap between axonal membrane and Schwann cell was around 30-40 nm (Fig. 2J). In addition, the distribution of TR-D3 in more periaxonal space were shown in spinal (Fig. S2C) and sciatic nerves (Fig. S2D) and plexus hepatis (Fig. S2E). Thus, the red TR-D3 fluorescence images co-stained with green sox-10 were also repeated in the spinal (Fig. S2F) and sciatic nerves (Fig. S2G) of wild type mice.

In the liver, axons labeled with cyan sox-10 are in a close proximity to the hepatic stellate cells labeled with green α-SMA (Fig. 2K). The red fluorescent TR-D3 therefore is capable to reach the extracellular space, possibly through axonal-stellate cells junctions as schematically depicted on Fig. 2L. EM images also support that the association between hepatic stellate cells and Schwann cells (Fig. 2M). Without the injection of fluorescence tracer, the observed immunofluorescence of green sox-10 with red α-SMA had concomitant relationship in liver (Fig. S2I)3

### Serotonin receptors in ependymal glia

Ventricular ependymocytes receive synaptic contacts and express receptors to various neurotransmitters ^18,19^; in particular these ependymocytes are innervated by serotonergic projections from raphe nucleus and express serotonin (5-HT) receptors ^20,21^. Whether ependymocytes of the central canal receive synaptic inputs were not previously studied. We show that the Thy1-YFP green fluorescent axons appear in close vicinity of ependymocytes stained by α-SMA (Fig 3A), the observed EM images also support the existence of axons and synapses around ependymocytes (Fig. 3B), although we cannot identify the neurotransmitter nature of these contacts.

**Figure 3.**
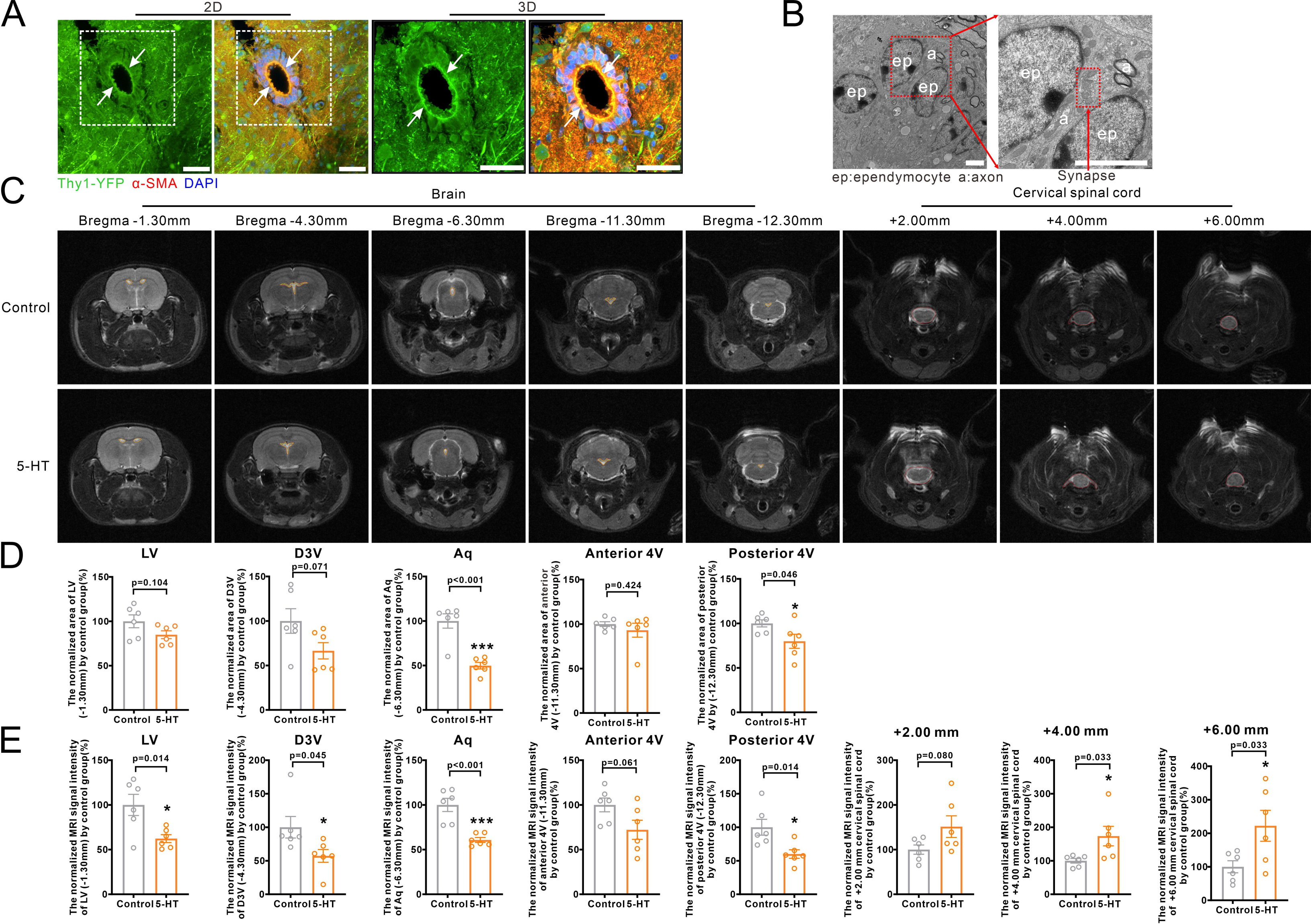
5-HT modulates the CSF distribution as revealed by MRI. A: Representative 2D and 3D images of spinal central canal surrounded with green YFP-labeled axons and co-stained with α-SMA (red) and DAPI (blue) in Thy1-YPF mice. Scale bar=50 μm. B: Representative electron microscope images of ependymocytes (ep), axon (a) and perisynaptic structure delineated by the box. Scale bar=2 μm. C: Representative MRI images of cerebral ventricles and cervical spinal cord in the rats treated with ACSF (control group) and in the rats treated with 5-HT. Yellow dotted lines delineate regions used to estimate the area of cerebral ventricles, and the red dotted lines delineate regions for MRI signal intensity measurements indicating the CSF content. D: The area of cerebral ventricles (labeled by yellow dotted lines) normalized by control group. Open circles represent data from individual rats (n=6) and bars represent mean±SEM, as comparison with control group *p<0.05, ***p<0.001. E: The MRI signal intensities of cerebral ventricles and cervical spinal cord (labeled by red dotted lines) normalized by control group. Open circles represent data from individual rats (n=6) and bars represent mean±SEM, as comparison with control group *p<0.05, ***p<0.001.

To test for serotonergic input we injected 5-HT at 8.8 ng/ml diluted in ACSF into CIM of rat, or 0.88 ng/ml into CIM of mice that corresponds to a final concentration of 5-HT in CSF ∼ 100 pmol/ml, which is within the physiological range of 1-100 pmol/ml ^22,23^. The T2-weighted MRI images were acquired from the brain and cervical spinal cord segments (Fig. 3C). Stronger signal intensity reflects more CSF (Fig. 3D and Fig. 3E). Image slices were separately collected at Bregma −1.30 mm, −4.30 mm, −6.30 mm, −11.30 mm and −12.3 mm, the corresponding area and signal intensities were calculated for lateral ventricle (LV), dorsal third ventricle (D3V), aqueduct (Aq), anterior and posterior fourth ventricle (4V). Ventricular area of LV, D3V and anterior 4V were not significantly changed after 5-HT injection (n=6; Fig. 3D). The area of Aq and posterior 4V were, however, significantly decreased by 5-HT (p<0.001 and p=0.046, n=6; Fig. 3D). Injection of 5-HT significantly decreased the CSF content in LV (p=0.014; n=6), D3V (p=0.045, n=6), Aq (p<0.001, n=6) and posterior 4V (p=0.014, n=6), but not in anterior 4V (p=0.061, n=6; Fig. 3E), as shown in Fig. 3C. Conversely, 5-HT significantly increased the MRI signal intensities in the cervical segments of the spinal cord (Fig. 3E). At +4.00 mm and +6.00 mm of cervical spinal cord, the signal intensities were significantly increased to 173.63% and 222.35% of control group (p=0.033, p=0.033; n=6). We also compared the MRI images before and after the CIM injection of 5-HT, as compared with the absence of 5-HT in the same rat. It turned out that 5-HT decreased the area and MRI signal intensities of cerebral ventricles, but increased the MRI signal intensities in cervical section of spinal cord (Fig. S3).

Expression of 5-HT_2B_Rs on ependymocytes in the central canal was confirmed with immunocytochemistry (Fig. 4A and Fig. S4A-F), which agrees well with the mRNA profile of 5-HT receptor subtypes in primary cultured ependymocytes ^24^. The 5-HT_2B_Rs were localized on both luminal and basal surface of ependymocytes, whereas F-actin was predominantly localized at the luminal surface with overlap between 5-HT_2B_Rs and F-actin. In the lateral ventricle, 5-HT_2B_R were localized to ependymocytes co-immunolabeled with α-SMA (Fig. S4G) or F-actin (Fig. S4H).

**Figure 4.**
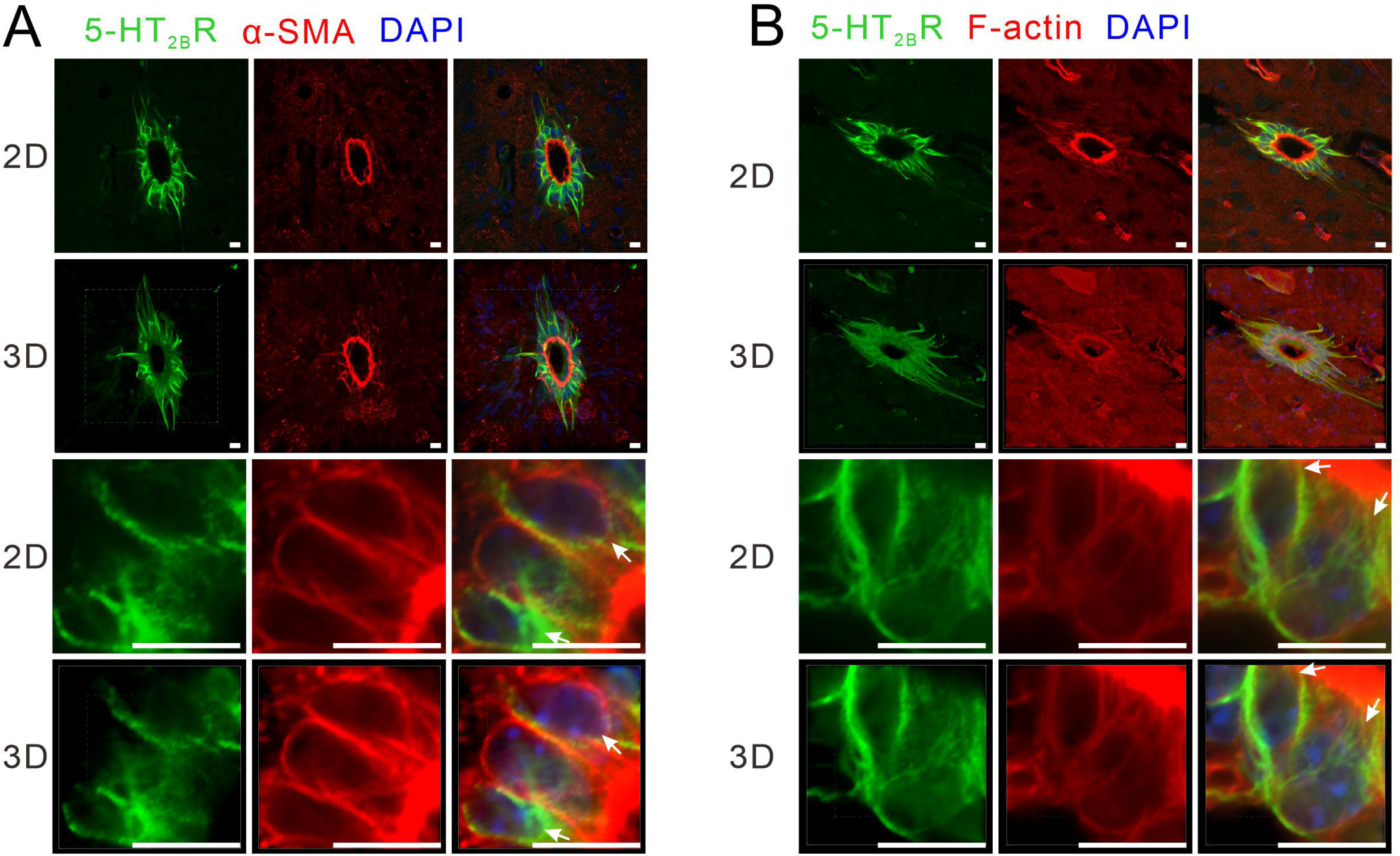
The expression of 5-HT_2B_R on ependymal cells of the central canal and lateral ventricle. A: 2D and 3D images of (green) canal co-stained with α-SMA (red) and DAPI (blue). Scale bar=10 μm. B: 2D and 3D images of co-stained with F-actin (red) and DAPI (blue). Scale bar=10μm.

### The effects of 5-HT on the ependymocytes

Functional consequences of stimulation of 5-HT receptors were first analyzed in primary cultured ependymocytes labeled with 5 μM calcein (Fig. 5A, 5B ad 5C). Addition of 5-HT to the culture medium at the final concentration of 100 nM initiated gradual decrease in the area of ependymocytes (Fig. 5B). In 12 minutes after 5-HT administration, the average cellular area decreased to 56.69% ± 1.62% (p<0.001, n=12) of the control values (Fig. 5C). Meanwhile, we detected the capacity of cellular recovery, 5-HT was added at 2 minutes in the flowing culture medium, the shrunk ependymocytes can be almostly recovered the initial area using 8.17 minutes (Fig. 5D-F; supplementary video 2). The recovered ependymocytes has no significant with the baseline after 10 minutes and 10 seconds (Fig. 5F). The pre-treatment with non-selective antagonist of 5-HT receptors, methylergometrine maleate (MM), as well as with a specific antagonist of 5-HT_2B_Rs, SB204741, prevented area decrease in response to 5-HT. In the concomitant presence of SB204741 and 5-HT, the area of ependymocytes was 77.09% ± 2.37% (p<0.001, n=12) of the control values (Fig. 5C). Next, to simulate their multi-cellular microenvironment *in vitro*, ependymocytes and astrocytes were co-cultured in the glass cubicles (Fig. S5A, S5B); again administration of 5-HT led to the contraction of ependymocytes (Fig. S5C, S5D). *In vivo,* changes in ependymocytes area translated into 5-HT-induced decrease of the area of the central canal (Fig. S5E and Fig. S5F) due to an increased flow of CSF into the spinal cord parenchyma. The area of the central canal was reduced following CIM injection of 5-HT (0.88 ng/ml diluted in ACSF) in mice, while the pre-treatment with either MM or SB204741 prevented the volume changes induced by 5-HT (Fig. S5G).

**Figure 5.**
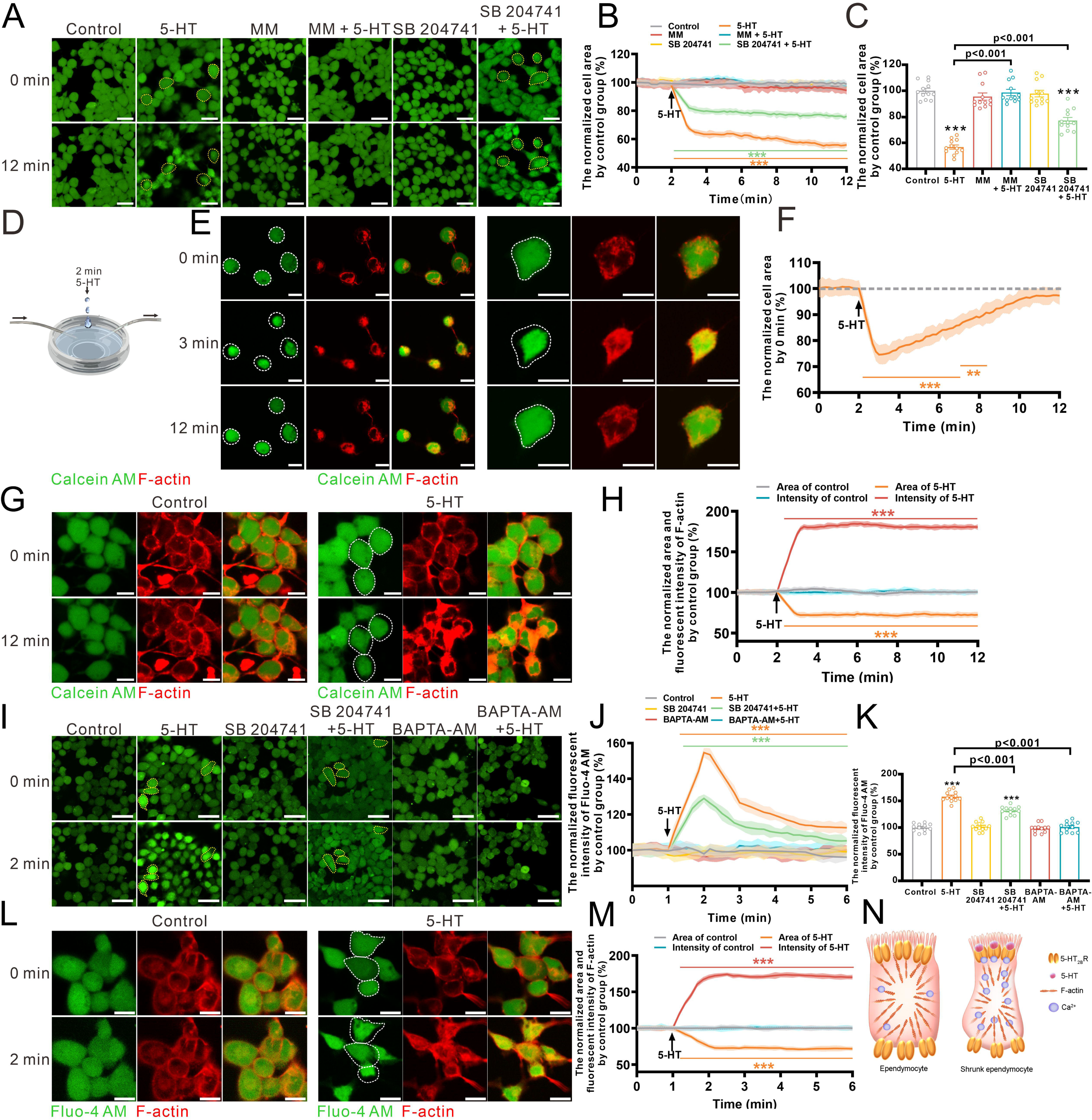
The 5-HT-induced ependymal shrinkage is mediated by 5-HT_2B_R and cytoplasmic Ca^2+^ increase induced F-actin polymerization. A; Representative image of calcein-AM labeled ependymocytes at 0 and 12 minutes of time lapse recording. Primary cultured cells were exposed (for 10 min) to 5-HT alone or after 15 min pre-treatment with methylergometrine maleate (MM; broad antagonist of 5-HT receptors) or SB204741 (selective antagonist of 5-HT_2B_R), then exposed to isotonic solution (control) or 5-HT for 10 minutes. Scale bar=25 μm. B: Changes of cellular area in response to 5-HT alone or in the presence of 5-HT receptors antagonists as indicated on the graph. Cell area was normalized to control group, every line shows mean values; while shadow shows SEM for every time point (acquired every 10 seconds); n = 12 experiments. ***p<0.001. C: The percentage of calcein-AM fluorescent intensity normalized by control group at 12 minutes. Open circles represent data from individual cultured batch (n=12) and bars represent averages±SEM, as comparison with control group ***p<0.001. D: Schematic of the added 5-HT into the flowing culture medium at 2 minutes. E: Representative images of calcein-AM labeled ependymocytes at 0, 3 and 12 minutes of time lapse recording. Primary cultured cells were exposed to 5-HT at 2 minutes and the culture medium was timely flowing. Scale bar=10 μm. F: Changes of cellular area in response to 5-HT. Cell area was normalized to control group, every line shows mean values; while shadow shows SEM for every time point (acquired every 10 seconds); n = 12 experiments. **p<0.01, ***p<0.001. G: Representative images of the calcein-AM and F-actin labeled ependymocytes. Primary cultured ependymocytes were labeled by calcein-AM and Sir-actin (F-actin), then incubated with 5-HT for 10 minutes. Scale bar=10 μm. H: Area and fluorescent intensity of F-actin normalized by control group at 12^th^ minute. Solid lines represent the mean for sepaarte cultures (n=12); shaded areas indicate SEM as compared to the control group ***p<0.001. I: Representative image of Fluo4-AM labeled ependymocytes in them presence of 5-HT alone or after 15 min pre-treatment with SB204741 or BAPTA-AM at 0 and 2 minutes of recording (peak of [Ca^2+^]_i_ increase). Scale bar=25 μm. J: Intracellular Ca^2+^ dynamics (expressed as Fluo-4 intensities normalized to control). Lines represent the mean±SEM for every time point (acquired every 10 seconds); n = 12 experiments. ***p<0.001. K: Fluo-4 fluorescent intensity normalized to control group at 2 minutes. Open circles represent means from individual (n=12) and bars represent averages±SEM, as compared to control group ***p<0.001. L: Representative images of the Fluo-4-AM and F-actin labeled ependymocytes incubated with 5-HT for 5 minutes. Scale bar=10 μm. M: Area and fluorescent intensity of F-actin normalized to control group at 6 minutes. Solid lines represent the means of individual experiments (n=12) and shaded areas indicate SEM, as compared to control group ***p<0.001. N: Schematic of the shrunk ependymocytes following polymerization of F-actin triggered by the 5-HT-induced [Ca^2+^]_i_ increase.

Changes in ependymocytes volume can be mediated through F-actin, the cytoskeletal protein regulating cell morphology and organelle mobility ^25^. We confirmed expression of F-actin in ependymocytes both *in vivo* and *in vitro*. Immunofluorescence *in situ* localized F-actin in the luminal surface of the ependymocytes co-stained for vimentin or α-SMA (Fig. S5H). The co-localization of F-actin with vimentin or α-SMA was also observed in the immunostained cultured ependymocytes (Fig. S5I). In live cultured ependymocytes, F-actin was also visualized with red fluorescence probe Sir-actin (Fig. 5G). Administration of 5-HT induced an areal change in ependymocytes associating with the polymerization and contraction of F-actin, as fluorescence intensity of F-actin increased together with decreased cellular surface (Fig. 5H).

Exposure of primary cultured ependymocytes to 5-HT significantly increased [Ca^2+^]_i_ (Fig. 5I and 5J left panel; supplementary video 3), to 157.90% ± 2.78% of the baseline (p<0.001; n=12) (Fig. 5K). Pre-treatment of cultures with SB204741 or with Ca^2+^ chelator BAPTA-AM suppressed 5-HT-induced [Ca^2+^]_i_ increase (n = 12) (Fig. 5I-K). When monitoring [Ca^2+^]_i_ together with F-actin, we found that 5-HT-induced increase in [Ca^2+^]_i_ was associated with the aggregation of F-actin (Fig. 5L and 5M). Treatment of cultures with BAPTA-AM prevented shrinkage of ependymocytes in response to 5-HT (Fig. S5J). Furthermore, the shrunk mechanism of ependymocytes induced by the [Ca^2+^]_i_ dependent F-actin aggregation was shown in Fig. 5N.

### Ependymocytes shrinkage induced by 5-HT affects CSF delivery to liver

Intra-CIM administration of 5-HT together with TR-D3 significantly increased the fluorescence of TR-D3 in hepatic stellate cells (p<0.001, n=6), while pretreatment with SB204741 (p=0.024, n=6) or BAPTA-AM (p=0.040, n=6) negated this effect (Fig. 6A-C). To further characterize the CSF movement from the ventricular system to the liver, the small molecular weight trisodium citrate dihydrate (C_6_H_5_Na_3_O_7_; MW=258.07) was injected into CIM at 70 mM diluted in 5 μl ACSF (18 mg/ml). After 4 hours, the brain, brain stem, thoracic portion of the spinal cord, hepatic plexus and liver were extracted, and the level of citrate in these tissues was measured by liquid chromatography mass spectrometry (LC-MS). The mass spectrum of the identified citrate and the standard concentration curve of trisodium citrate dihydrate are shown in Fig. S6A and Fig. S6B, and the value of trisodium citrate dihydrate normalized by protein content is shown in Fig. S6C and Fig. S6D. As compared with the control levels from same tissues (Fig. 6D), 5-HT significantly decreased citrate levels by 28.82% ± 3.64% in the brain (p<0.001, n=6) and by 39.15% ± 4.91% in the brain stem (p<0.001, n=6), whereas the citrate level was increased by 32.72% ± 5.14% in the thoracic spinal cord (p<0.001, n=6), by 39.36% ± 5.19%, in hepatic plexus (p<0.001, n=6) and by 60.35% ± 5.97% in the liver (p<0.001, n=6). The pre-treatment with SB204741 and BAPTA-AM prevented 5-HT effects on the citrate propagation.

**Figure 6.**
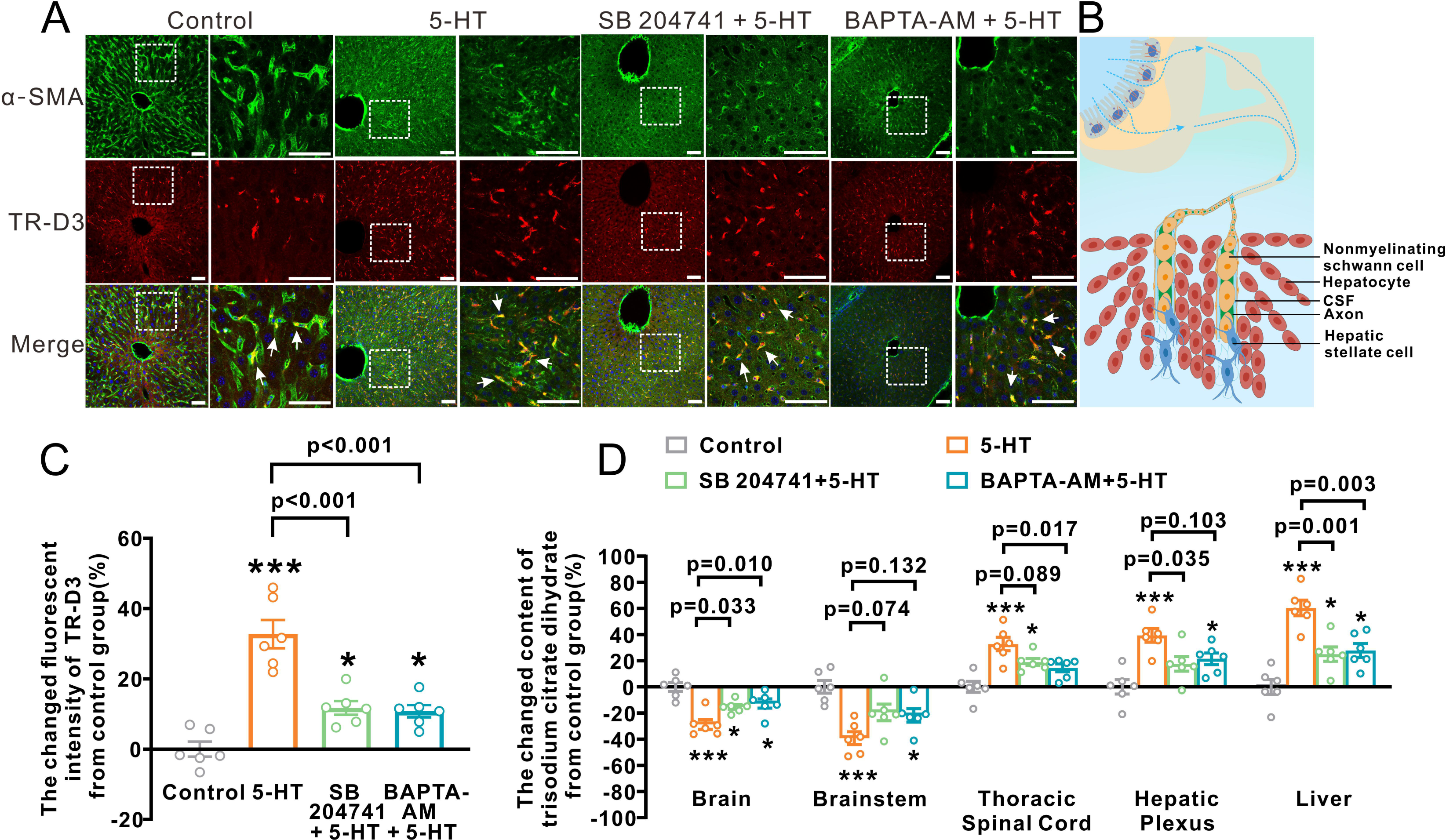
Delivery of fluorescent markers form the CNS into liver is 5-HT_2B_R and Ca^2+^ dependent. A: Representative images of TR-D3 fluorescence co-stained with α-SMA (green) and DAPI (blue) in the liver. Scale bar=100 μm. B: Schematic depiction of CSF flow through periaxonal space (PAS) into liver. C: Fluorescence intensities of TR-D3 normalized to control; open circles represent data from individual mice (n=6) and bars represent mean±SEM, *p<0.05, **p<0.01. D: Trisodium citrate dihydrate levels measured by HPLC-MS 4 hours after CIM injection in the brain, brain stem, thoracic spinal cord, hepatic plexus and liver. Open circles represent data from individual mice (n=6) and bars represent averages±SEM, as comparison with control group *p<0.05, **p<0.01, ***p<0.001.

Subsequently, we measured fluorescence tracers (TR-D3 and FITC-D40) in acutely isolated brain slices from the forebrain, brain stem, the cervical, thoracic and lumbar spinal cord segments and in the liver measured at 4 hours after CIM injection (Fig. 7 and Fig. S7). Exposure to 5-HT decreased fluorescence intensity of FITC-D40 in the anterior and posterior brain, by 61.62% ± 1.39% (p<0.001, n=6) and 50.77% ± 1.23% (p<0.001, n=6) of the control, whereas TR-D3 fluorescence was reduced by 56.92% ± 3.19% (p<0.001, n=6) and 56.93% ± 5.56% (p<0.001, n=6). In the brain stem, the intensities of FITC-D40 and TR-D3 decreased by 61.78% ± 1.71% (p<0.001, n=6) and 54.36% ± 2.91% (p<0.001, n=6) of the control group, respectively. In the cervical portion of the spinal cord, the intensities of FITC-D40 and TR-D3 were increased by 68.26% ± 3.16% (p<0.001, n=6) and 56.30% ± 3.93% (p<0.001, n=6) of control group. Similar tendency was observed in slices from thoracic and lumbar parts of the spinal cord. Intensities of FITC-D40 were elevated by 40.88% ± 3.61% (p<0.001, n=6) and 96.37% ± 3.72% (p<0.001, n=6) of control group, while the intensities of TR-D3 were increased by 70.20% ± 2.70% (p<0.001, n=6) and 71.12% ± 4.27% (p<0.001, n=6) of control group. Pretreatment with MM, SB204741 or BAPTA-AM fully antagonized effects of 5-HT on FITC-DC and TR-D3 propagation from CNS to liver (Fig. 7C and S7B).

**Figure 7.**
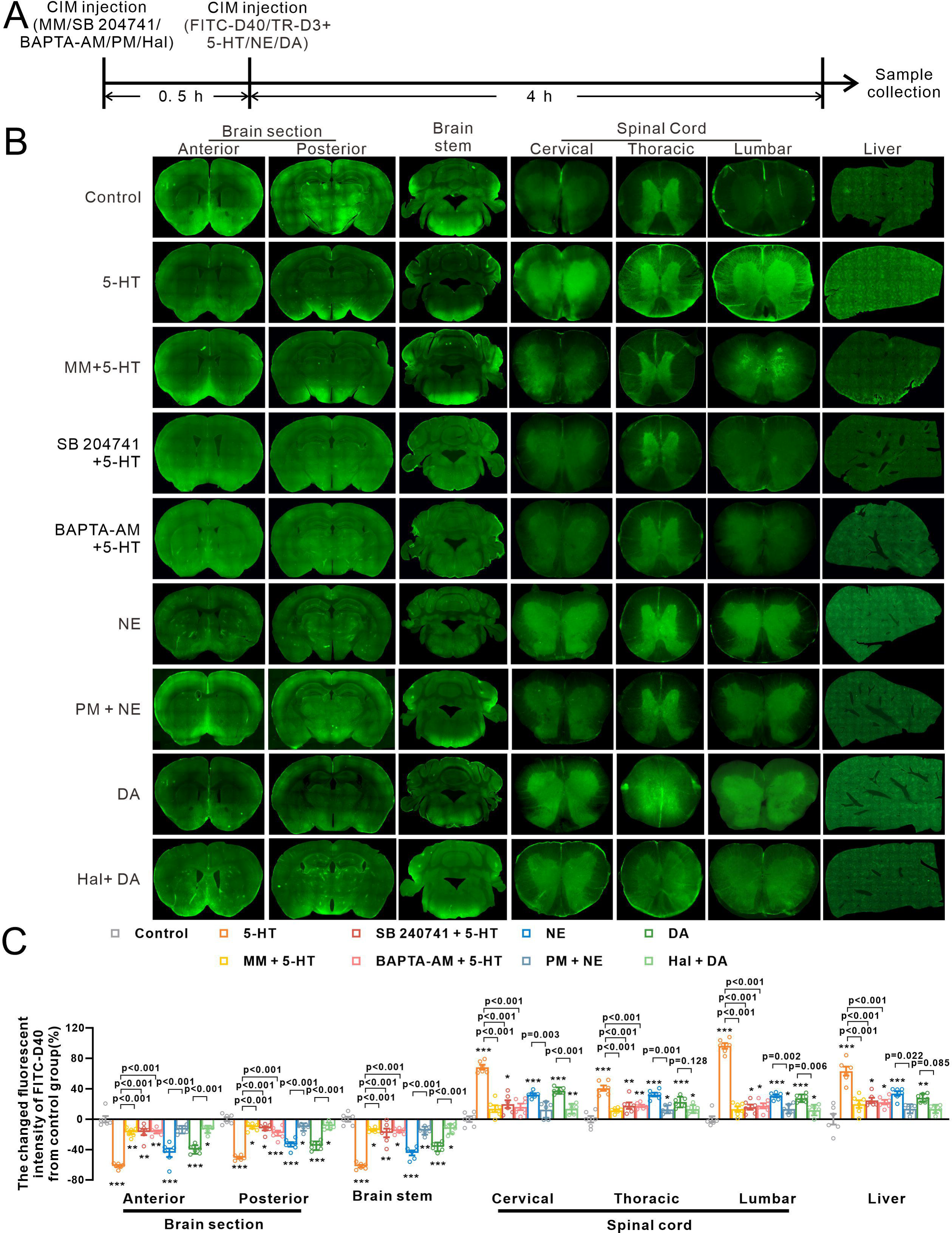
Delivery of CSF-derived FITC-D40 to the liver is regulated by catecholamines. A: The time schedule of CIM injection. B: Representative images of fluorescence of FITC-D40 in the anterior and posterior brain sections, brainstem, the cervical, thoracic and lumbar spinal cord, and liver in control and in the presence of 5-HT, norepinephrine (NE), and dopamine (DA). C: Fluorescence intensities of FITC-D40 in different tissues in the presence of catecholamines and respective antagonists of relevant receptors. Open circles represent data from individual mice (n=6) and bars represent mean±SEM, *p<0.05, **p<0.01, ***p<0.001.

### Noradrenaline and dopamine increase CSF delivery to the liver

Serotonin was not the only neurotransmitter affecting CSF movement from the CNS to peripheral organs. Spinal cord ependymocytes potentially express other catecholamine receptors, as indeed ependymocytes in 3^rd^ ventricles were shown to express adrenoreceptors as well as receptors to dopamine ^26^. Norepinephrine (NE) and dopamine (DA) both increased the delivery of fluorescence tracers to the spinal cord and liver, and decreased their presence in the brain and brain stem (Fig. 7B and S7B). Inhibition of adrenoceptors by phentolamine mesylate effectively prevented the NE-induced increase in CSF propagation. Likewise haloperidol, an inhibitor of dopamine receptors antagonized DA effects on CSF delivery from CNS to liver (Fig. 7C and S7B).

## Discussion

Here we studied in detail the connection between the CSF and peripheral organs through the peripheral nerves. We found that tracers with low (cadaverine, 1 kDa) medium (TD-3R, 3 kDa) and high molecular weight (FITC-D40) after being injected into CIM distribute first trough the spinal cord and then are delivered to the liver and pancreas through the peripheral nerves. We further demonstrated that spinal cord ependymocytes can regulate CSF movement from the spinal canal toward peripheral organs by active volumetric changes, this gating CSF flow along peripheral nerves periaxonal route (PAS). This discovery advances the previous knowledge about the CSF movement, and shows that neurotransmitter-induced changes in the volume of ependymal cells can control the one-way flow of CSF from central canal through spinal cord parenchyma into PAS. Volume changes in ependymocytes regulated by neurotransmitters are mediated through Ca^2+^-dependent polymerization and contraction of cytoskeletal F-actin. Serotonin-evoked shrinkage of ependymocytes decreased the presence of CSF in cerebrum while increasing CSF in the spinal cord and liver. In the hepatic interstitium, the connection between Schwann cells and hepatic stellate cells facilitates CSF entry into the liver parenchyma.

### Ependymocytes regulate CSF movement

Ependymal glia form the wall of the ventricular system and the central canal, erecting a partial barrier between CSF and CNS parenchyma ^11,27^. Ventricular ependymocytes receive synaptic catecholaminergic innervation from brainstem nuclei ^19,21^. Here we present the data indicating likely innervation of ependymocytes of the central canal of the spinal cord; we further demonstrate functional expression of 5-HT_2B_ receptors in the spinal cord ependymal cells (Fig. 4, S4A-F, 5A and S5G). Ependymocytes belong to astroglia, and hence possess classical intracellular excitability, mainly mediated by Ca^2+^ signaling ^28^, which is triggered by neurochemical stimulation. Ependymal Ca^2+^ signals were characterized in response to 2 mM glucose ^29^, as well as by activation of P2X_7_ purinoceptors ^30,31^. Here we demonstrate that stimulation of metabotropic 5HT_2B_receptors triggers Ca^2+^ signaling in ependymal cells; this Ca^2+^ signaling in turn induces polymerization of F-actin and volume change of ependymocytes. It is conceivable that under physiological conditions, the monoamines regulate the rhythmical fluctuations of CSF flow mediated by dynamic change in the volume of ependymal cells. It is also likely that ependymocytes at different parts of the ventricles and central canal have distinct sensitivity to neurotransmitters that increase the versatility of CSF flow regulation.

In the central canal, the ependymocytes possess high levels of cytoskeleton protein F-actin (Fig. 4B and Fig. S5H-S5I), which may mechanistically contribute to the changes in the ependymocytes shape and volume ^32,33^. In addition, F-actin network is involved in coordinating cilia movement at the apical surface ^34^. Cytosolic Ca^2+^ signals regulate F-actin ^35^, in particular Ca^2+^ binds to F-actin to form Ca^2+^-F-actin ^36^. Polymerization of F-actin in *vibrio splendidus* agglutinated by a subunit of oligomerization domain-like receptors (NLRs) is Ca^2+^-dependent ^37^. Similarly, cytoplasmic Ca^2+^ regulates the reorganization of F-actin in BeWo cell line ^38^. We propose that stimulation of 5-HT_2B_Rs triggers Ca^2+^ signals that interact with F-actin to dynamically control the shape and volume of ependymocytes. Resulting shrinkage of ependymocytes opens intercellular space for CSF flow into the spinal parenchyma in one-way direction towards peripheral nerves. Ependymocytes work as the water-gates, which can be regulated by synaptic input and control the CSF flow out of the lumen of the central canal to the spinal cord parenchyma and further to the periaxonal space of spinal nerves.

### CSF flow through peripheral nerves periaxonal route

It is well known that CSF clears metabolic waste produced by the brain parenchyma, while the CSF flow is rhythmically fluctuating ^39^. It is generally accepted that ependymal cell cilia beat rhythmically to assist the CSF flow, and CSF mainly circulates in the parenchyma of brain and spinal cord, the cerebral ventricles and central canal. The CSF outflow along the peripheral spinal nerves was hypothesized ^9,10^ but was not directly proven. In this study, we demonstrate that ependymocytes act as dynamic regulators of CSF flow from the spinal cord central canal into the spinal cord parenchyma and subsequently to the periaxonal space of peripheral nerves, by which CSF can reach peripheral organs. Opening of the ependymal gate by catecholaminergic neurotransmitters (5-HT, NE and DA) acting through receptors expressed in ependymocytes actively increases the redistribution of CSF from the brain to the spinal cord and then to peripheral organs (Fig. 7 and Fig. S7). Thus, the CSF-mediated exchange between the CNS and the periphery represents the tightly regulated pathway for information and material exchange. Both metabolites and signaling molecules can use this pathway to coordinate peripheral functions with overall activity of the brain and the spinal cord.

### Conclusions and future perspectives

We found that ependymocytes of the central canal of the spinal cord can dynamically and rapidly change their shape and volume under the action of monoamines. Ependymocytes morphological plasticity controls the CSF flow from the central canal to the peripheral nerves and by proxy to the peripheral organs. The CSF flow may be instrumental in delivery of various signaling molecules and metabolites from the CNS to the periphery, thus adding another level of complexity of CNS-dependent control of visceral functions. In this paradigm, the CSF acts as a signaling medium extending from the brain core to the peripheral organs. Further investigations of this system and its regulation may be instrumental for understanding the physiology of CNS - periphery communications in health and disease.

### Limitations of the study

The ependymal regulation of CSF-based CNS-periphery signaling was characterized in rodents, and this needs to be validated and confirmed in humans. Direct monitoring of ependymal shrinkage in the central canal of live animals is hard to accomplish; hence, we characterized the changes of the central canal volume in the freshly isolated tissues of the spinal cord. In MRI results, the changes of the spinal central canal of rat was hard to be identified, hence we only analyzed the area of cerebral ventricles.

## Acknowledgments

This work was supported by the National Natural Science Foundation of China, MX [grant number 32271038] and BL [grant number 81871852]; Shenyang Young and Middle-aged Science and Technology Innovation Talent Support Program, BL [grant number RC210251]; Chunhui Project Foundation of the Education Department of China, BL [grant number 2020703].

## Authors contributions

A.V., M.X. and B.L. designed and supervised the study. X.L., D.Z., S.W., Y.F., Y.L. and W.Y. collected the data *in vitro* and analysed the relevant data. X.L., D.Z., S.W., S.L., and L.C. collected the data *in vivo* and analysed the relevant data. B.L. M.X. and A.V. wrote the manuscript and B.L., M.X., A.V. and T.H. edited it.

## STAR Methods

### Key resources table

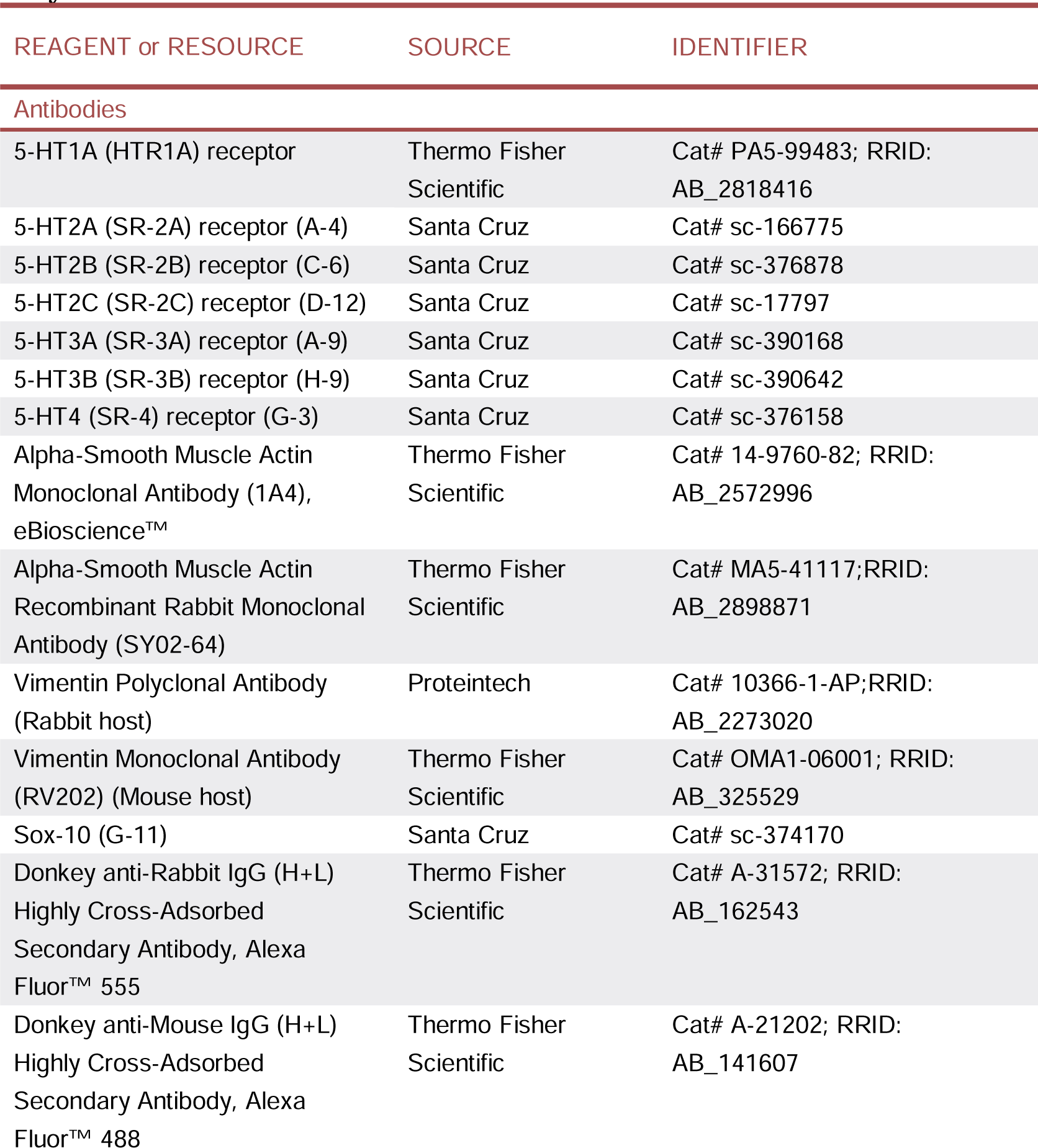
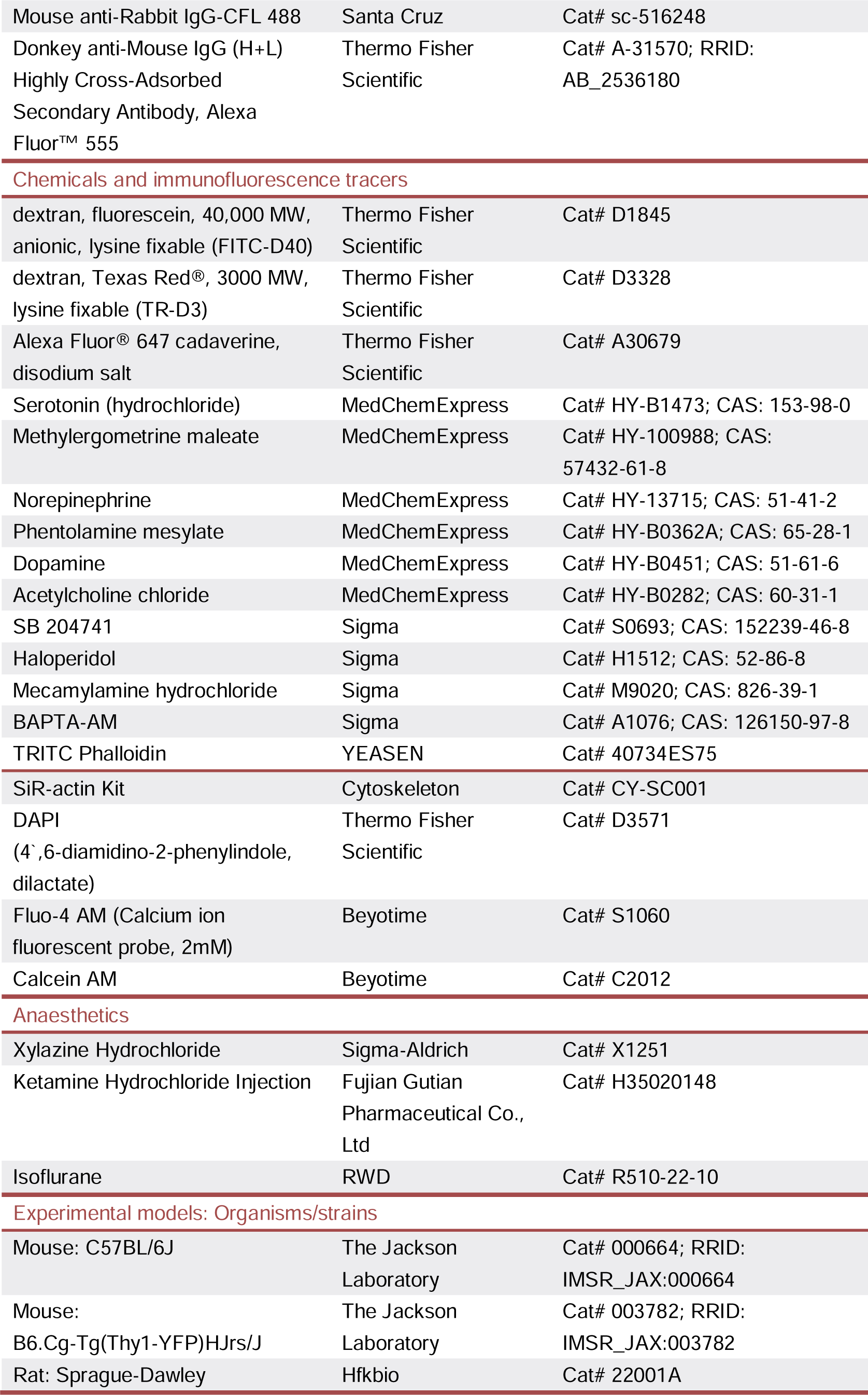
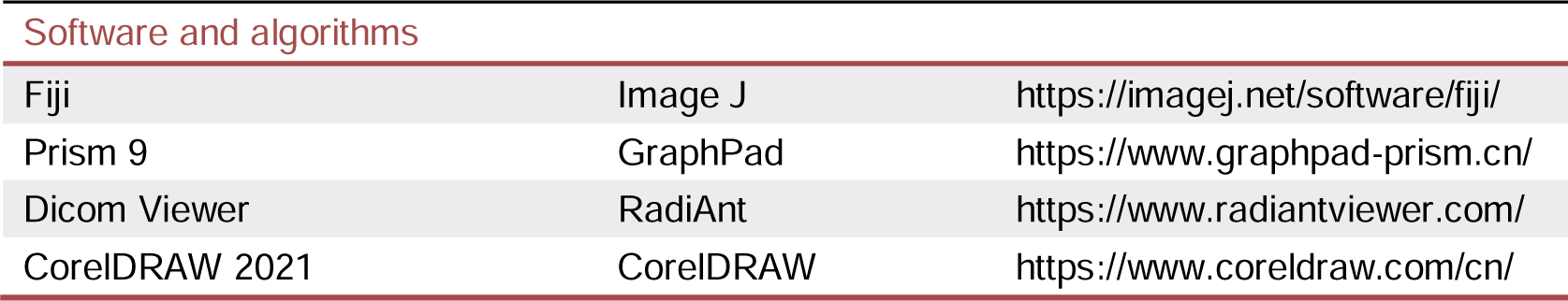

### Resource Availability

### Lead Contact

Further information and requests for resources and reagents should be directed to and will be fulfilled by the lead contact, Baoman Li.

### Materials Availability

This study did not generate any new reagents.

### Data and code availability

Source data and statistical analysis underlying the main and supplementary figures are available in Supplementary Data.

### Experimental model and subject details Animals

The male wild-type C57BL/6J mice (#000664) and B6.Cg-Tg(Thy1-YFP)HJrs/J (#003782) transgenic mice were purchased from the Jackson Laboratory (Bar Harbor, ME, USA). Male Sprague-Dawley rats (#22001A, purchased from Hfkbio, Beijing, China) were used for MRI experiments. The mice and rats were kept in standard rooms at 22 ± 1 °C, maintained on a 12/12 h light/dark cycle, and were given adequate food and water. All experiments were performed under the guidance of the US National Institutes of Health Guide for the Care and Use of Laboratory Animals (NIH Publication No. 8023) and its 1978 revision. All experimental protocols were approved by the Institutional Animal Care and Use Committee of China Medical University, approval No. [2020]102.

## Method details

### Tracers Injection and Detection

We monitored distribution of fluorescent tracers between different parts of the CNS and peripheral organs to trail CSF movements ^40–42^. The fluorescent tracers TR-D3 and FITC-D40 were dissolved in artificial cerebrospinal fluid (ACSF) at 1 % (w/v). The male adult mice were fasted before surgery, and anesthetized using ketamine (100 mg/kg) and xylazine (10 mg/kg). Head stereotaxic system was used to fix mouse while surgically exposing the posterior atlanto-occipital membrane. The tracers were delivered to CSF through cisterna magna injection using a 30 GA needle, at a rate of 1 μl/min for 5 minutes (5 μl total volume). In 4 hours after tracer injection, anesthetized mice were fixed by transcardially perfusion with phosphate buffer saline (PBS) and 4 % paraformaldehyde (PFA). Tissues were cut into 50 μm slices, images captured using Carl Zeiss Axio Scan microscope (Promenade 10, Jena, Germany) or confocal scanning microscope (DMi8, Leica, Wetzlar, Germany).

ImageJ software (ImageJ 1.46r, NIH) was used to calculate and analyze fluorescent images. Intensities of FITC-D40 and TR-D3 from different groups were normalized to the intensity of the control group. For images scanned with Carl Zeiss AxioScan microscope, we used ImageJ to select regions of interests to calculate mean fluorescence intensity of FITC-D40 and TR-D3; ImageJ was also used to remove background color and quantify the mean fluorescence intensity of TR-D3.

### Immunofluorescence Assay and Calculation

As described previously ^40^, the anaesthetized male adult mice were perfused through the heart with PBS and 4 % PFA, soaked in 4 % PFA and kept for 24 hours. After collection, the tissues were cut into uniform slices with a thickness of 60 μm. Tissue slices were incubated with donkey serum for 1 h for permeabilization. Primary antibodies were incubated at 4 °C overnight, followed by secondary antibody incubated at room temperature for 2 hours. The staining of F-actin is tissue slices incubation with 100 nM TRITC-phalloidin (YEASEN Biotechnology, Shanghai, China) for 30 minutes at room temperature after secondary antibodies incubation. Nuclei of cells were marked with its common immunofluorescence marker, 4’, 6’-diamidino-2-phenylindole (DAPI) at 1:1000 dilution. Images were captured using a confocal scanning microscope (DMi8, Leica, Wetzlar, Germany). The specific information on the antibodies used is shown in the table.

### Ligation of vena cava and sectioning of the spinal nerves

Adult male mice were anesthetized with ketamine (100 mg/kg) and xylazine (10 mg/kg). To ligate the head branch of the superior vena cava, the chest cavity was carefully opened to expose the heart, then the head branch of the superior vena cava was ligated. To disconnect the spinal nerves, the mice were anesthetized as above, bilateral erector spinal muscles were separated to expose the spinal nerves from T2 to T12 on both sides; the nerves were then cut. After the surgical operations, the mice were quickly sutured and kept body temperature around 37 °C until the full awakening.

### Tissue Sampling

The anaesthetized mice were perfused and fixed with PBS and 4 % PFA through the heart. After 15 minutes, the mouse brain, brain stem, spinal cord, liver, pancreas, cervical lymph nodes, spinal nerve roots and sciatic nerve were taken for subsequent immunofluorescence experiments. The spinal nerve root is taken from the cervical segment of the spinal cord, located after the intersection of the anterior and posterior roots (near the intervertebral foramen).

### Two-photon Microscopy

10 to 12-week-old male B6.Cg-Tg(Thy1-YFP)HJrs/J transgenic mice were anesthetized with ketamine (100 mg/kg, i.p.) and xylazine (10 mg/kg, i.p.), mice body temperature was kept around 37 °C. The spinal nerves at thoracic level from T7-T9 were exposed, and the observation window of the exposed spinal nerves was perfused with ACSF and observed under the objective. The images of the nerve were taken every three seconds using FluoView with two-photon laser-scanning setup (Nikon AR1, Japan). Bandpass filters (Chroma) were 540nm/40nm for YFP and 850nm/70nm for TR-D3 signals. The confocal images of the nerve were taken every 3 seconds. The red tracer TR-D3 was injected into CIM at a rate of 1 μl/min for 5 minutes (5 μl total volume). In 35 – 55 min after injection the tracer appeared in the spinal nerves, the observations were started after the injection of TR-D3, the first presence of tracer TR-D3 under observation view was set as 0 s.

### Electron Microscopy

In order to fix the liver, spine, and sciatic nerve of mice, phosphate-buffered saline (PBS) and 2.5% glutaraldehyde were combined. Dissected samples of the liver, spine, and sciatic nerve were cut into 1 mm3 pieces and fixed at 4°C in 2.5% glutaraldehyde. After being fixated in a solution consisting of 1% osmium for two hours at 4°C, the pieces were rinsed multiple times in PBS, dried on a graduated series of ethanol concentrations ranging from 20% to 100%, then using 100% acetone, penetrated by Epon 812, and then polymers on pure Epon 812 for 72 hours at 65°C. The samples were set up in comparatively thin portions. Using an ultramicrotome, extremely thin slices (70 nm) were cut, gathered in copper grids, and painted using lead citrate and uranyl acetate. The samples in every component underwent TEM (JEM-1400Flash, Japan) examination in 10 visible areas.

### MRI analysis

Rats were anesthetized with 3% and 2% isoflurane. Head stereotaxic instrument was used to fix rats while surgically exposing the posterior atlantooccipital membrane. The ACSF (Control group) or serotonin (5-HT group) was delivered into the CSF of rat through cisterna magna injection at a rate of 1 μl/min for 5 minutes using a 30 GA needle. The injected rats were sutured and then placed in the MRI animal imaging system (Bruker biospec3T, Bruker, Billerica, MA USA), T2-weighted images were scanned and image acquisition was performed using ParaVision software (ParaVision PV-360.2.0.p1.1, Bruker, Billerica, MA USA). The scan parameters for the T2-weighted images sequence were: axial scan direction, repetition time=4538 ms, effective TE=85 ms, echo time=15 ms, flip angle=90°, number of averages=8, field of view=35 × 35 mm^2^, matrix size=256×256, slice thickness=1 mm, number of slices=25. ImageJ (ImageJ 1.46r, NIH) was used to delineate and calculate the areas and MRI signal intensities of the lateral ventricle, aqueduct, and fourth ventricle, as well as the MRI signal intensities of the high signal area at the cervical spinal cord. The calculated ventricle area of the selected optical planes (yellow dotted lines) was normalized by the total area of the brain tissue in the same optical planes, in order to exclude the individual difference. The liquid content of the selected tissues in the optical planes (red dotted lines) was reflected by MRI signal intensity. At last, the measured data in every group were all normalized to the control group.

### Ependymal Cells Culture

Primary cultured ependymal cells were prepared based on the experimental operation described previously ^43^. The newborn mouse brains were aseptically isolated, the tissue was vortexed, filtered and suspended in MEM_C_ (MEM containing 5 mg/l insulin, 0.5 g/l BSA and 10 mg/l transferrin), the suspensions were planted in fibronectin-treated cell culture dishes. After forty-eight hours of static culture, changed the culture medium to MEMcT (MEMc with 500 U/l thrombin). Refresh the medium completely every three days.

### Co-Culture of Ependymocytes with Astrocytes in Glass Cubicles

Newborn mice within 3 days of birth were used to extract astrocytes as reported previously ^42,44^. The neopallia of cerebral hemispheres were isolated and made into single-cell suspensions under sterile conditions. Isolated astrocytes were cultured in a moist environment of carbon dioxide / air at 37 ℃, using Dulbecco Minimum Essential Medium (DMEM), with 15% fetal bovine serum as culture medium.

As described previously ^45^, the cells were harvested (using 0.125 % trypsin) and seeded in fibronectin-coated 24 well Transwell^®^ chamber insert (Hirao et al., 2023). The upper chambers were seeded with ependymal cells, and the lower chambers were seeded with astrocytes. Follow-up experiments were carried out when the degree of cell confluence reached about 80 %.

### Cell Volume Assay

For cell volume monitoring and imaging in cultured ependymocytes, cells were treated with 3 μM calcein AM (Beyotime, Shanghai, China) for 15 minutes. After PBS containing calcein AM was replaced with PBS without calcein AM, cells were incubated for 10 minutes, and then tests were carried out. The cell volume of cultured ependymocytes were visualized by calcein signals at 10-s intervals for 12 minutes using confocal scanning microscope (DMi8, Leica, Wetzlar, Germany). The cell volume from all calcein positive cells in a field of recording selected of every culture dish used in experiment were included in the statistics, the cell volume of ependymocytes was normalized to the baseline volume in 0 minute. The test was repeated in 12 different cultures using the same method. ImageJ software (ImageJ 1.46r, NIH) was used to quantify the volume.

### Cellular Recovery Detection

Cultured ependymocytes were treated with 1 μM Sir-Actin (Cytoskeleton, Denver, CO, USA) for 30 minutes. After PBS containing Sir-Actin was replaced with PBS without Sir-Actin, cells were incubated with calcein AM as above. Images of cultured ependymocytes co-treated with calcein AM and Sir-Actin were captured at 10-s intervals for 12 minutes using confocal scanning microscope (DMi8, Leica, Wetzlar, Germany). During the imaging, two identical constant-flow pumps (BT100-1J, Longer, Hebei, China) were used to input and output the culture medium at 40 rpm (∼100 μL/min) to keep it flowing and to maintain complete equilibrium of the liquid level. The cell volume from all calcein positive cells in a field of recording selected of every culture dish used in experiment were included in the statistics, the cell volume of ependymocytes was normalized to the baseline volume in 0 minute. The test was repeated in 12 different cultures using the same method. ImageJ software (ImageJ 1.46r, NIH) was used to quantify the volume.

### Ca^2+^ Imaging and Quantification

As described previously ^46^, cultured ependymocytes were incubated with 3 μM fluo-4 AM (Beyotime, Shanghai, China) for 15 minutes. Subsequently PBS containing Fluo-4 was replaced with PBS without Fluo-4, in which cells were incubated for 10 minutes. The fluorescence signals of Fluo-4 were monitored at 10-s intervals for 6 minutes by confocal scanning microscope (DMi8, Leica, Wetzlar, Germany). As previously described ^46^, all cells labeled with Fluo-4 in a field of recording were included in the statistics. Fluo-4 intensity was normalized to the baseline intensity at the beginning of experiment. The ImageJ software (ImageJ 1.46r, NIH) was used to analyze Ca^2+^ signals. The measurements were repeated in 12 different cultures.

### F-actin Assay

To monitor and image F-actin in ependymocytes, cells were treated with 1 μM Sir-Actin (Cytoskeleton, Denver, CO, USA) for 30 minutes. After PBS containing Sir-Actin was replaced with PBS without Sir-Actin, cells were incubated with calcein AM or fluo-4 AM as above. Images of cultured ependymocytes co-treated with Sir-Actin and calcein AM were captured at 10-s intervals for 12 minutes while that of co-treated with Sir-Actin and fluo-4 AM were captured at 10-s intervals for 6 minutes using confocal scanning microscope (DMi8, Leica, Wetzlar, Germany). The intensity of F-actin area from all Sir-Actin positive cells in a field of recording were included in the statistics, the intensity of F-actin was normalized to the baseline area or intensity at 0 minute. Both tests were repeated in 12 different cultures. ImageJ software (ImageJ 1.46r, NIH) eas used to analyze all images.

### Tissue Volume Detection

Mice were sacrificed by decapitation, and the spinal cord was removed immediately. The cervical spinal cord was cut into 500 μm slices and incubated with 5 μM calcein AM (Beyotime, Shanghai, China) dissolved in phosphate buffer saline (PBS) for 15 minutes. After PBS containing calcein AM was replaced with PBS without calcein AM, tissues were incubated for 10 minutes before recordings. Images were captured at 10-s intervals for 12 minutes by using a confocal scanning microscope (DMi8, Leica, Wetzlar, Germany). The cross-sectional area of central canal was normalized to the baseline area in 0 minute. Experiment was repeated in 6 separate preparations. Images were analysed with ImageJ sowtware (ImageJ 1.46r, NIH).

### High Performance Liquid Chromatograph Mass Spectrometer (HPLC-MS)

The trisodium citrate dihydrate were diluted in ACSF at a concentration of 18 mg/ml. The anaesthetized mice were injected with trisodium citrate dihydrate into the cisterna magna at a rate of 1 μl/min for 5 minutes (5 μl total volume), and 4 hours later, the mice were decapitated. 30 mg each of the mice brain tissue, brain stem, thoracic spinal cord and liver, and 5 mg of the hepatic plexus were homogenized at 4 °C and 60 Hz for 3 minutes using tissue grinder (Scientz-48L, SCIENTZ, Ningbo, China). The tissue homogenate was dissolved with Methanol: acetonitrile: water 1:1:2 (v/v/v) solution, centrifuged at 13,000 rpm, and take the supernatant for detection by LC-MS.

The liquid chromatography system (1260 Infinity, Agilent, Santa Clara, USA) was used to separate the test substance before mass spectrometry. The autosampler was set up at room temperature, and the temperature of column was kept at 30 °C. The flow rate of mobile phase is kept at a uniform rate of 0.5 mL/min, and the volume of injection of all samples is always kept at 10 μL. Chromatographic separation was accomplished on a InfinityLab Poroshell 120 SB-C18 column (100 × 4.6 mm i.d., 2.7 μm; Agilent, Santa Clara, USA), the testing time is unified as 3 min. Mobile phase (acetonitrile: water = 70:30, v/v, 0.1% formic acid) were freshly prepared. To prevent for potential carryover, the ratio of methanol to water is maintained at 1: 1, v/v.

A mass spectrometer (6420 Triple Quad LC/MS, Agilent, Santa Clara, USA) equipped with a negative turbo ionspray electrospray ionization source was used. Parameters for mass spectrum detection were set as described below: ion-spray voltage, 3500 V; gas temperature, 310 °C. Using protonated molecule [M-H]-ion for all analyzes. The quantifier and qualifier process used twin multiple reaction monitoring (MRM) transitions. The quantitative MRM was set at mass-to-charge ratio (m/z) 191.0 → 111.0. Stock solutions of trisodium citrate dihydrate were prepared using methanol to final concentrations of 200 mg/ml. Working trisodium citrate dihydrate solutions were freshly produced before every batch analysis, were prepared in Methanol:acetonitrile:water = 1:1:2 (v/v/v) at concentration of 10, 100, 1000, and 10000 μg/ml.

### Citrate quantification in tissues

Chromatography data were acquired and analyzed using Agilent MassHunter Workstation Software LC/MS Data Acquisition (version B.07.00, Agilent) and Qualitative Analysis (version B.06.00, Agilent). Using standard curve to calculate the citrate content in tissue samples, the content of citrate from different groups was normalized to the content of the control group.

### Statistical analysis

One-way ANOVA with Tukey post hoc test was used for comparisons included more than two groups, unpaired two-tailed t-test was used for two-group comparisons, GraphPad Prism 9 software (GraphPad Software Inc., La Jolla, CA, USA) and SPSS 24 software (International Business Machines Corp., NY, USA) were used for the above statistics and analyses. All statistical data in the text were presented as the mean ± SEM, the value of significance was set at p < 0.05.

---

**Figure S1.**
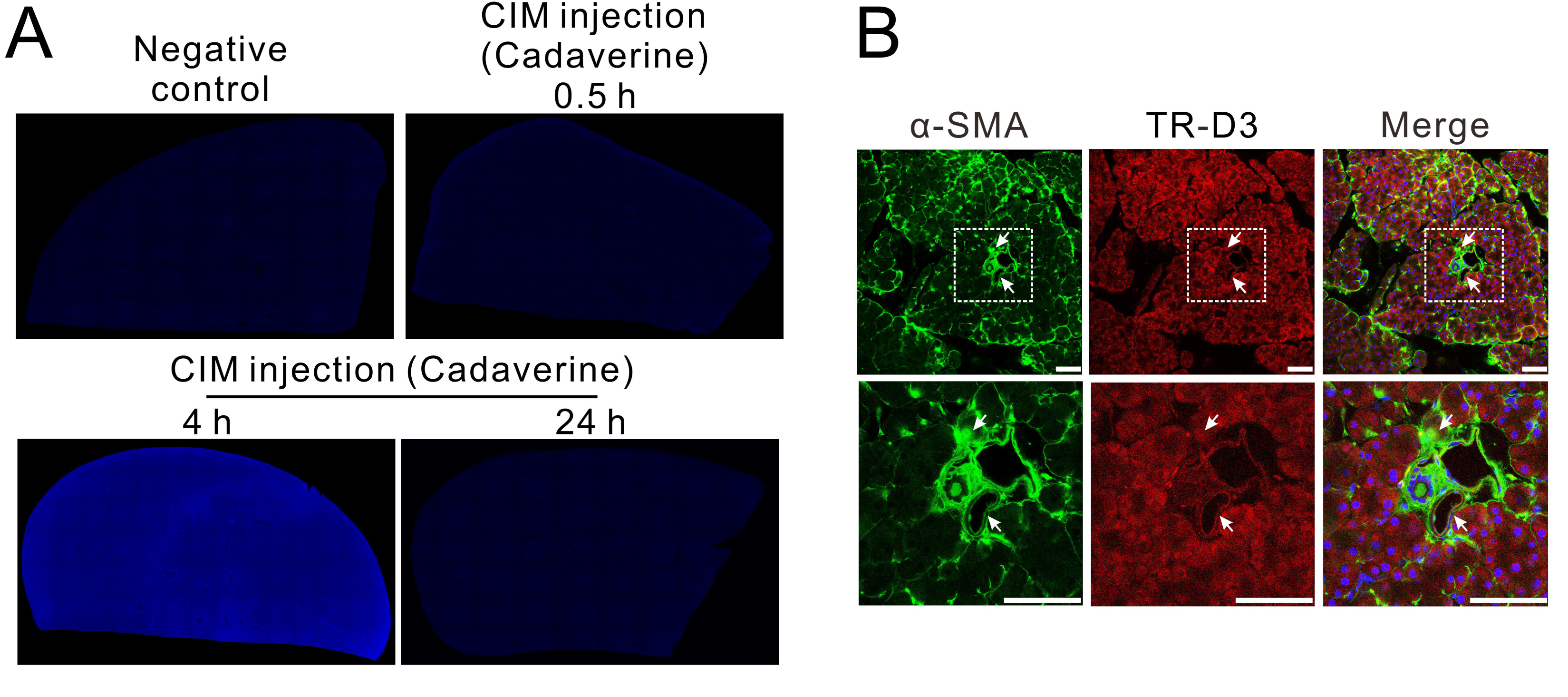
The peripheral distribution of tracers injected from cisterna magna. A; Representative fluorescence images of cadaverine in the liver 30 minutes, 4 hours and 24 hours after CIM injection with the negative control. B: Representative fluorescence images of TR-D3 (red) in pancreas co-stained with α-SMA (green) and nucleus marker DAPI (blue). Scale bar = 100 μm.

**Figure S2.**
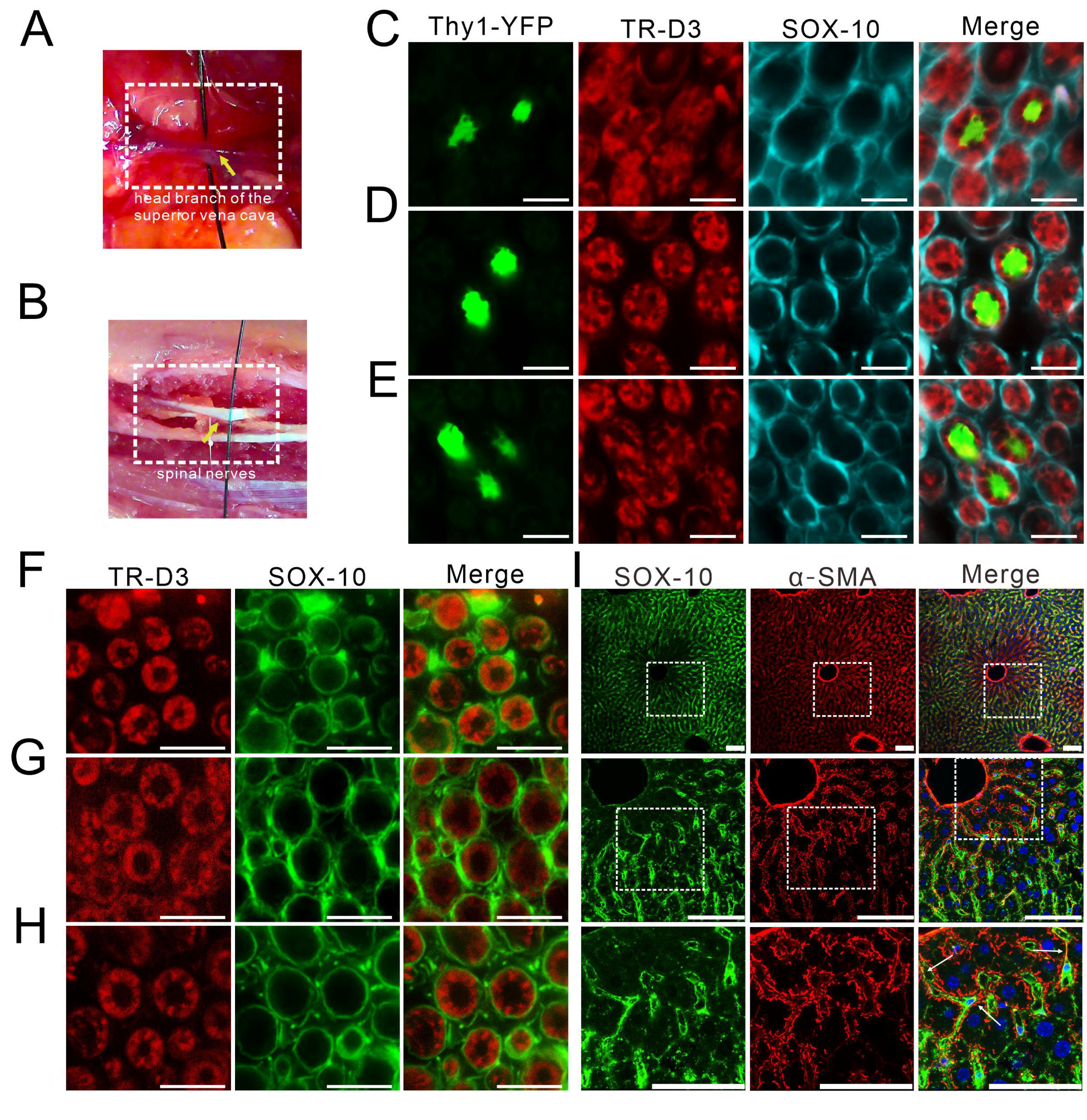
The peripheral periaxonal pathway of CSF. A; Ligated head branch of superior vena cava. B: Sectioned spinal nerves. C: Representative cross section images of fluorescence TR-D3 (red) sox-10 (cyan) and Thy1-YFP (green) in the periaxonal space of spinal nerves 4 hours after TR-D3 was injected into CIM. Scale bar=10 μm. D: Representative cross section images of fluorescence TR-D3 (red) sox-10 (cyan) and Thy1-YFP (green) in the periaxonal space of sciatic nerves 4 hours after TR-D3 was injected into CIM. Scale bar=10 μm. E: Representative cross section images of fluorescence TR-D3 (red) sox-10 (cyan) and Thy1-YFP (green) in the periaxonal space of the nerve roots separated from the hepatic nerve plexus 4 hours after TR-D3 was injected into CIM. Scale bar=10 μm. F: Representative cross section images of fluorescence TR-D3 (red) and sox-10 (cyan) in the periaxonal space of spinal nerves 4 hours after TR-D3 was injected into CIM of wild type mice. Scale bar=20 μm. G: Representative cross section images of fluorescence TR-D3 (red) and sox-10 (cyan) in the periaxonal space of sciatic nerves 4 hours after TR-D3 was injected into CIM of wild type mice. Scale bar=20 μm. H: Representative cross section images of fluorescence TR-D3 (red) and sox-10 (cyan) in the periaxonal space of the nerve roots separated from the hepatic nerve plexus 4 hours after TR-D3 was injected into CIM of wild type mice. Scale bar=20 μm. I: Representative images of fluorescence sox-10 (green) and α-SMA (red) immunofluorescence in the liver. Scale bar=100 μm.

**Figure S3.**
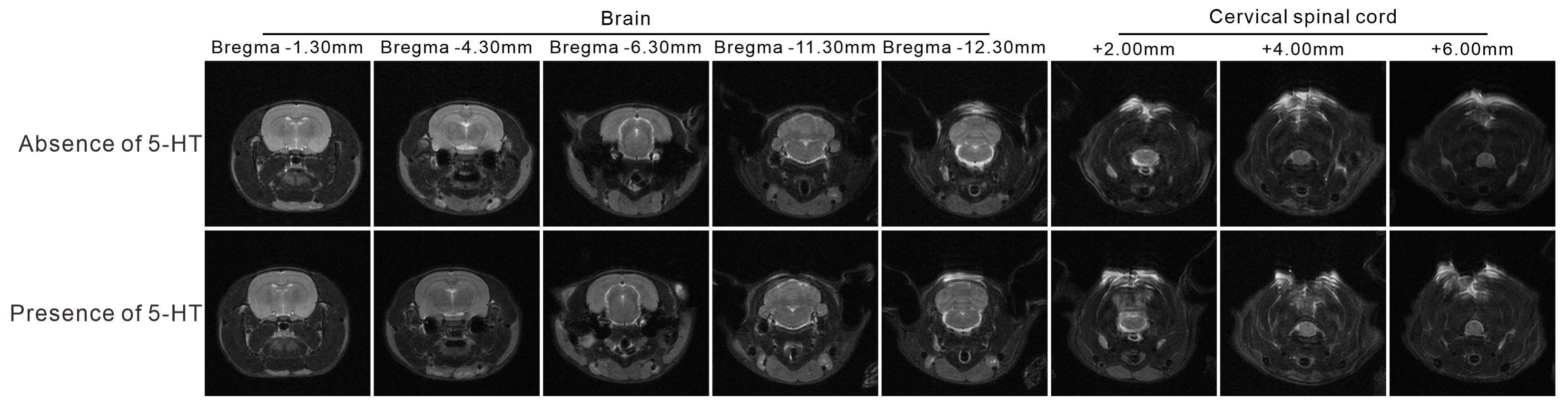
MRI images rat before and after CIM injection of 5-HT. Representative MRI images of the same rat in the absence (upper panels) and presence of 5-HT (lower panels).

**Figure S4.**
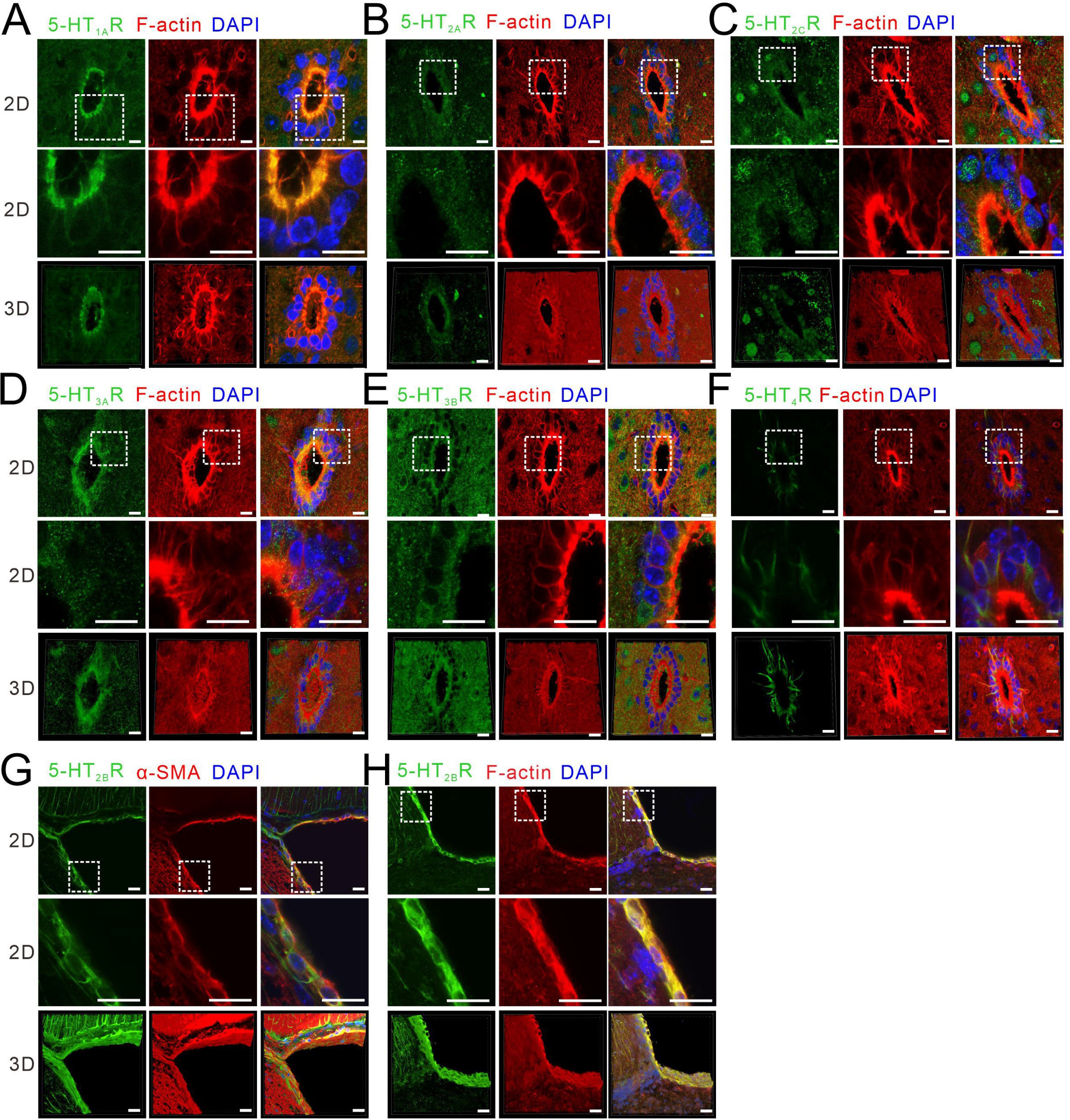
Molecular profile of 5-HT receptors expressed on the ependymocytes of the spinal central canal. A-F: 2D and 3D images of green 5-HT_1A_R (A), 5-HT_2A_R (B), 5-HT_2C_R (C), 5-HT_3A_R (D), 5-HT_3B_R (E) and 5-HT_4_R (F) in ependymal cells of spinal central canal co-stained with F-actin (red) and DAPI (blue), Scale bar=20 μm. G: 2D and 3D images of 5 co-stained with a-SMA (red) and DAPI (blue) on ependymal cells. Scale bar=20 μm. H: 2D and 3D images of-HT_2B_Rs (green) in ependymocytes of of lateral ventricle co-stained with F-actin (red) and DAPI (blue) on ependymal cells of lateral ventricle. Scale bar=20 μm.

**Figure S5.**
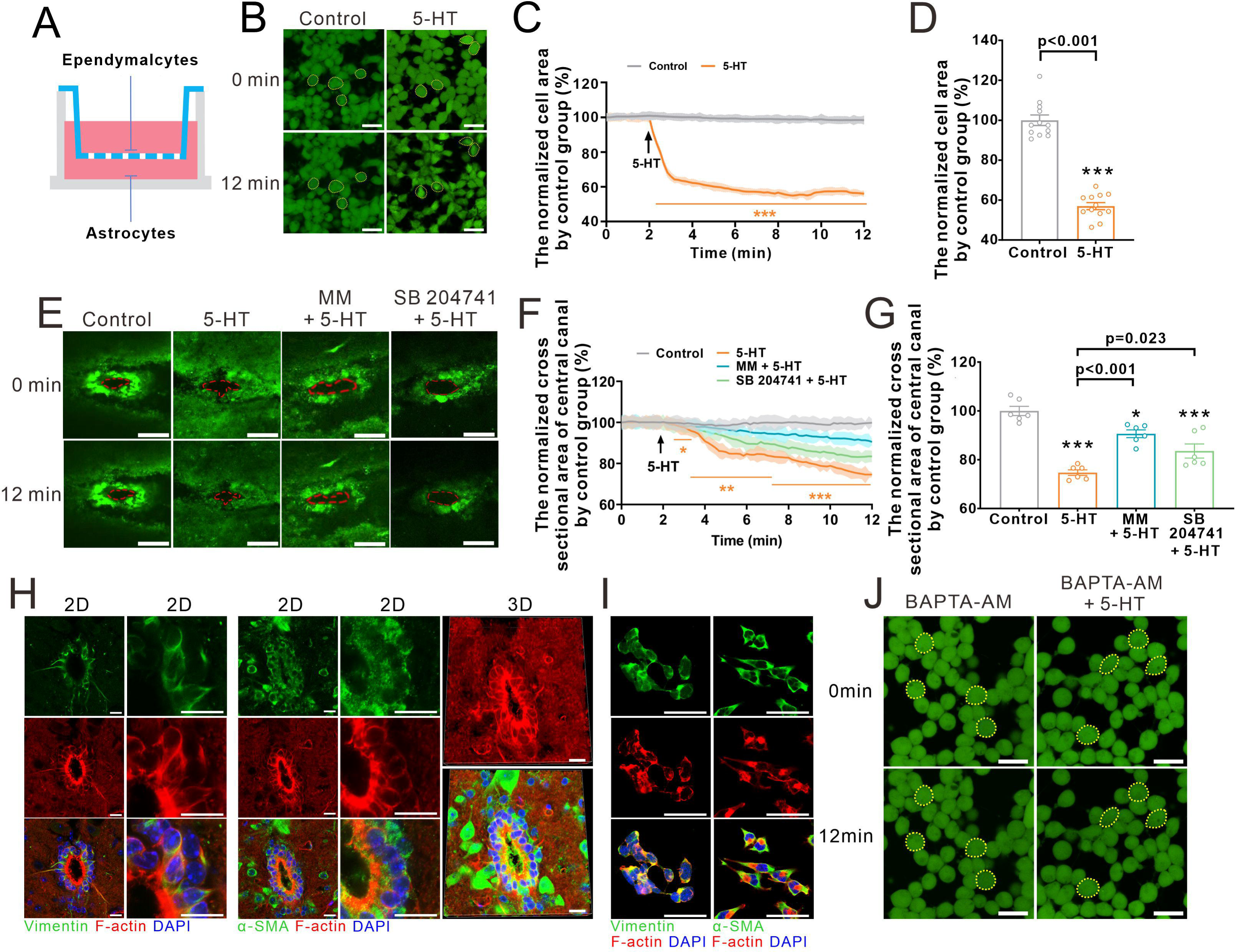
5-HT triggered ependymocytes shrinkage by stimulating Ca^2+^-induced F-actin polymerization in primary and co-cultured ependymocytes. A: The schematic of co-culturing ependymocytes with astrocytes in glass cabinets. B: the representative image of calcein-AM labeled ependymocytes treated with equilibration solution (control) or 5-HT at 0 and 12 minutes. Scale bar=25 μm. C: Time course of the changes in ependymocytes area.Solid lines represent the means of individual experiments (n=12) and shaded areas indicate SEM for every time point (acquired every 10 seconds), as compared to control group ***p<0.001. D: Cell area ratio at 12^th^ minute. Open circles represent data from every experiment (n=12); bars show means±SEM, as compared with control group ***p<0.001. E: Representative images of calcein-AM labeled acute slices of the spinal central canal at 0 and 12 minutes of time lapse recording. After the collection of fresh spinal cord tissues, the central canal in was pre-treated with MM or SB204741 for 15 minutes, then PBS (control) or PBS +5-HT were applied for 10 minutes. Scale bar=50 μm. F: Changes of the area of central canal in response to 5-HT alone or in the presence of 5-HT receptors antagonists as indicated on the graph. The data points were normalized to control values, every line represents mean±SEM for every time point (acquired every 10 seconds), n = 12 experiments. *p<0.05, **p<0.01, ***p<0.001. G: Central canal area normalized to control group at 12 minutes. Open circles represent data from individual cultured batch (n=12) and bars represent averages±SEM, as comparison with control group *p<0.05, ***p<0.001. H: Representative 2D and 3D images of F-actin (red) co-stained with vimentin (green) or α-SMA (green) and DAPI (blue) in spinal central canal. Scale bar=20 μm. I: Representative 2D images of F-actin (red) co-stained with vimentin (green) or α-SMA (green) and DAPI (blue) in primary cultured ependymocytes. Scale bar=20 μm. J: Representative images of calcein-AM labeled ependymocytes at 0 and 12 minutes. The primary cultured cells were pre-treated with BAPTA-AM for 15 minutes, then were treated with PBS (control) or PBS+5-HT for 10 minutes. Scale bar=25 μm.

**Figure S6.**
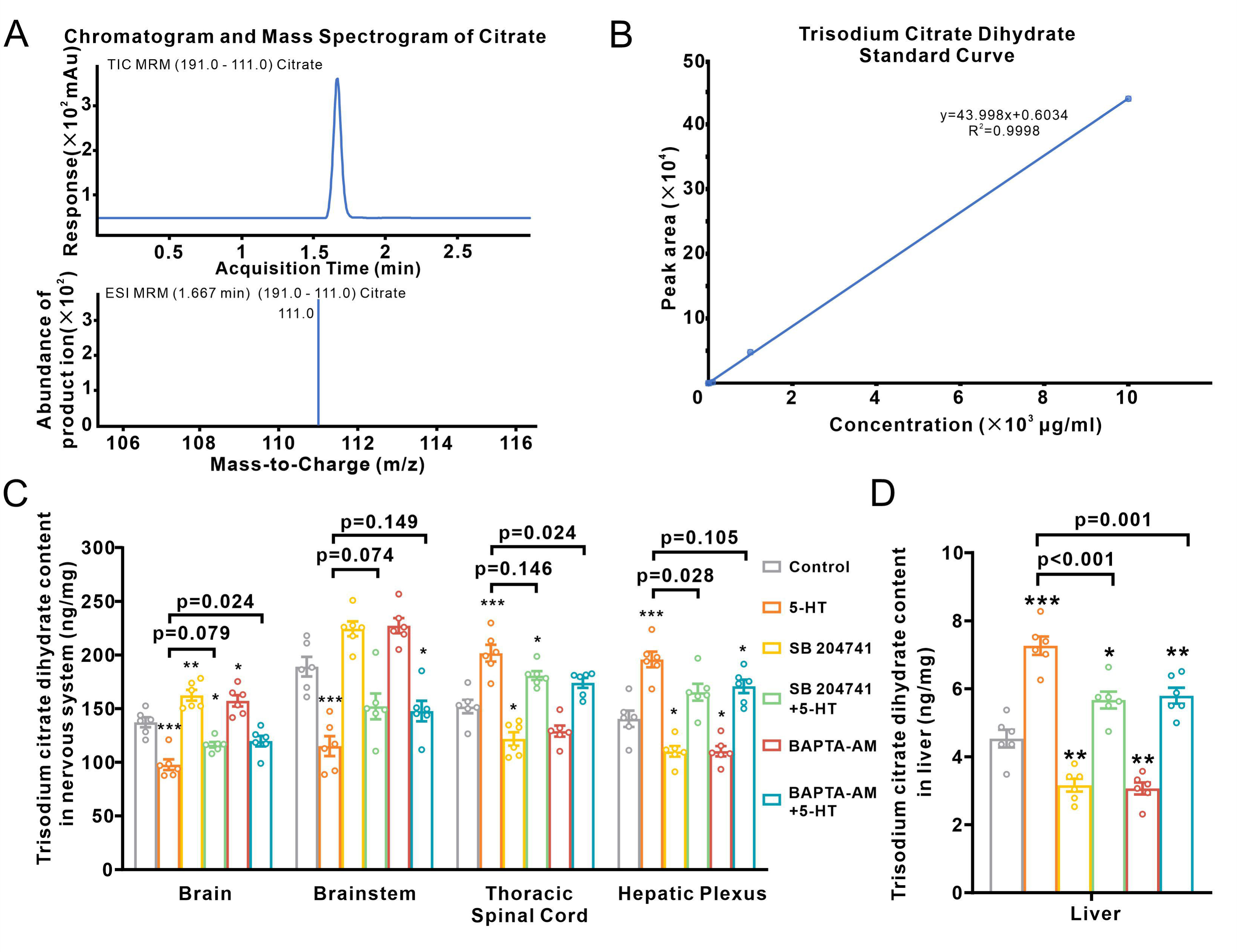
Distribution of trisodium citrate measured by HPLC-MS in different tissues. A: The mass spectrogram images of citrate in acquisition time (top) and mass to charge. B: The standard curve of measured citrate by HPLC-MS. C: The measured value of trisodium citrate dihydrate content normalized to the protein weight in brain, brain stem, thoracic spinal cord and hepatic plexus. Open circles represent data from individual mice (n=6) and bars represent averages±SEM, as comparison with control group *p<0.05, **p<0.01, ***p<0.001. D: Trisodium citrate dihydrate content normalized by the protein weight in liver. Open circles represent data from individual mice (n=6) and bars show mean±SEM, *p<0.05, **p<0.01.

**Figure S7.**
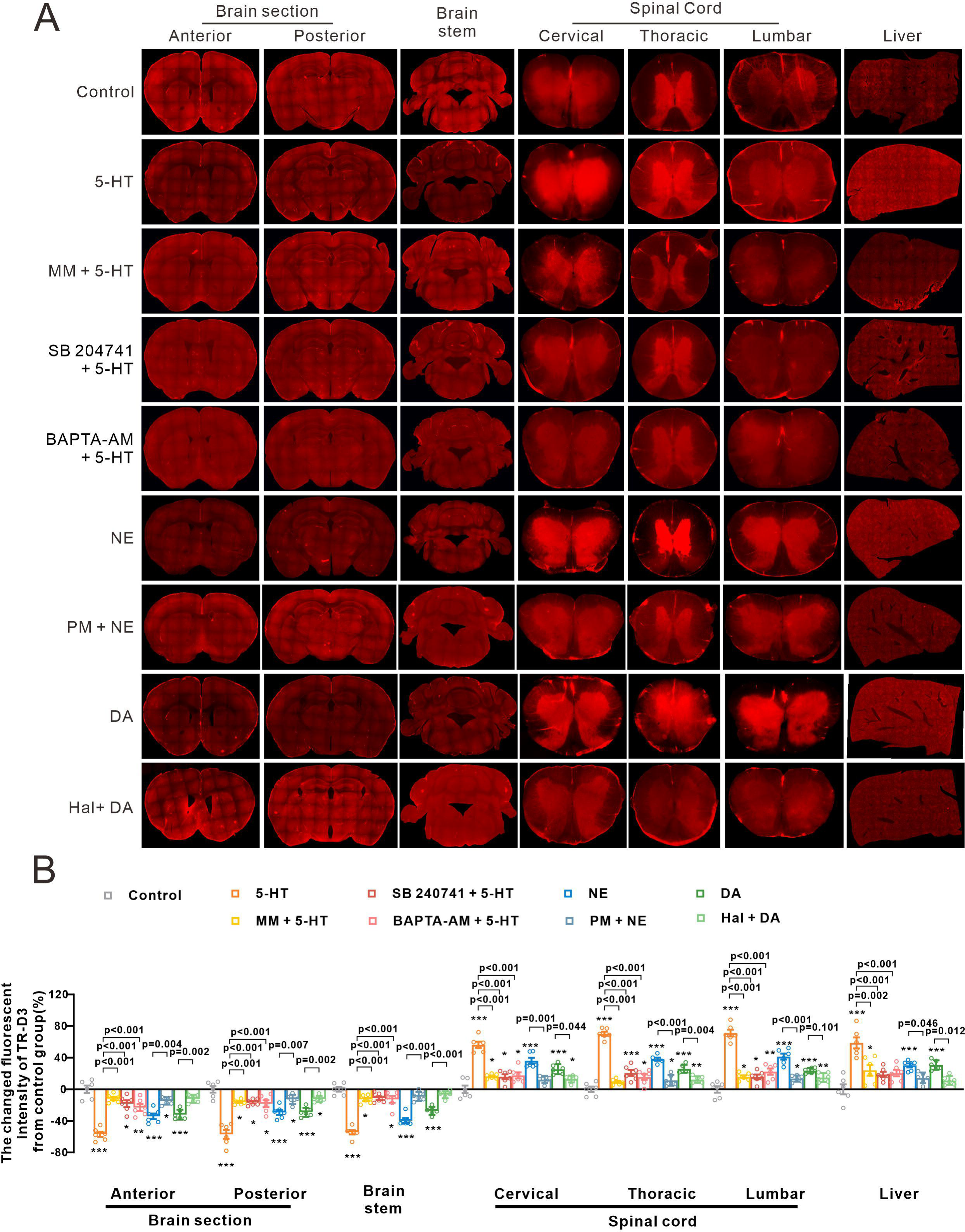
Delivery of CSF-derived TR-D3 to the liver is regulated by catecholamines. A: Representative images of fluorescence of TR-D3 in the anterior and posterior brain sections, brainstem, the cervical, thoracic and lumbar spinal cord, and liver in control and in the presence of 5-HT, norepinephrine (NE), and dopamine (DA). B: Fluorescence intensities of TR-D3 in different tissues in the presence of catecholamines and respective antagonists of relevant receptors. Open circles represent data from individual mice (n=6) and bars represent mean±SEM, *p<0.05, **p<0.01, ***p<0.001.

## References

1. Alperin, N., Bagci, A.M., Lee, S.H., and Lam, B.L. (2016). Automated Quantitation of Spinal CSF Volume and Measurement of Craniospinal CSF Redistribution following Lumbar Withdrawal in Idiopathic Intracranial Hypertension. AJNR. American journal of neuroradiology 37, 1957–1963. 10.3174/ajnr.A4837.

2. Brinker, T., Stopa, E., Morrison, J., and Klinge, P. (2014). A new look at cerebrospinal fluid circulation. Fluids Barriers CNS 11, 10. 10.1186/2045-8118-11-10.

3. Chazen, J.L., Dyke, J.P., Holt, R.W., Horky, L., Pauplis, R.A., Hesterman, J.Y., Mozley, D.P., and Verma, A. (2017). Automated segmentation of MR imaging to determine normative central nervous system cerebrospinal fluid volumes in healthy volunteers. Clin Imaging 43, 132–135. 10.1016/j.clinimag.2017.02.007.

4. Hodel, J., Lebret, A., Petit, E., Leclerc, X., Zins, M., Vignaud, A., Decq, P., and Rahmouni, A. (2013). Imaging of the entire cerebrospinal fluid volume with a multistation 3D SPACE MR sequence: feasibility study in patients with hydrocephalus. European radiology 23, 1450–1458. 10.1007/s00330-012-2732-7.

5. Spector, R., Robert Snodgrass, S., and Johanson, C.E. (2015). A balanced view of the cerebrospinal fluid composition and functions: Focus on adult humans. Exp Neurol 273, 57–68. 10.1016/j.expneurol.2015.07.027.

6. Wichmann, T.O., Damkier, H.H., and Pedersen, M. (2021). A Brief Overview of the Cerebrospinal Fluid System and Its Implications for Brain and Spinal Cord Diseases. Frontiers in human neuroscience 15, 737217. 10.3389/fnhum.2021.737217.

7. Hablitz, L.M., and Nedergaard, M. (2021). The Glymphatic System: A Novel Component of Fundamental Neurobiology. The Journal of neuroscience: the official journal of the Society for Neuroscience 41, 7698–7711. 10.1523/jneurosci.0619-21.2021.

8. Alpár, A., Benevento, M., Romanov, R.A., Hökfelt, T., and Harkany, T. (2019). Hypothalamic cell diversity: non-neuronal codes for long-distance volume transmission by neuropeptides. Current opinion in neurobiology 56, 16–23. 10.1016/j.conb.2018.10.012.

9. Bechter, K. (2011). The peripheral cerebrospinal fluid outflow pathway –physiology and pathophysiology of CSF recirculation: A review and hypothesis. Neurology, psychiatry and brainresearch 17,51–66.

10. Proulx, S.T. (2021). Cerebrospinal fluid outflow: a review of the historical and contemporary evidence for arachnoid villi, perineural routes, and dural lymphatics. Cellular and molecular life sciences: CMLS 78, 2429–2457. 10.1007/s00018-020-03706-5.

11. Verkhratsky, A. & Butt, A. (2023) Neuroglia: Function and pathology. Academic Press, Elsevier, pp 730.

12. Alpár, A., Zahola, P., Hanics, J., Hevesi, Z., Korchynska, S., Benevento, M., Pifl, C., Zachar, G., Perugini, J., Severi, I., et al. (2018). Hypothalamic CNTF volume transmission shapes cortical noradrenergic excitability upon acute stress. The EMBO journal 37. 10.15252/embj.2018100087.

13. Serra, R., and Simard, J.M. (2023). Adherens, tight, and gap junctions in ependymal cells: A systematic review of their contribution to CSF-brain barrier. Frontiers in neurology 14, 1092205. 10.3389/fneur.2023.1092205.

14. Li, Y.H., Woo, S.H., Choi, D.H., and Cho, E.H. (2015). Succinate causes α-SMA production through GPR91 activation in hepatic stellate cells. Biochem Biophys Res Commun 463, 853–858. 10.1016/j.bbrc.2015.06.023.

15. Li, J., Zeng, C., Zheng, B., Liu, C., Tang, M., Jiang, Y., Chang, Y., Song, W., Wang, Y., and Yang, C. (2018). HMGB1-induced autophagy facilitates hepatic stellate cells activation: a new pathway in liver fibrosis. Clin Sci (Lond) 132, 1645–1667. 10.1042/cs20180177.

16. Ahn, J.H., Cho, H., Kim, J.H., Kim, S.H., Ham, J.S., Park, I., Suh, S.H., Hong, S.P., Song, J.H., Hong, Y.K., et al. (2019). Meningeal lymphatic vessels at the skull base drain cerebrospinal fluid. Nature 572, 62–66. 10.1038/s41586-019-1419-5.

17. Pizzo, M.E., Wolak, D.J., Kumar, N.N., Brunette, E., Brunnquell, C.L., Hannocks, M.J., Abbott, N.J., Meyerand, M.E., Sorokin, L., Stanimirovic, D.B., and Thorne, R.G. (2018). Intrathecal antibody distribution in the rat brain: surface diffusion, perivascular transport and osmotic enhancement of delivery. J Physiol 596, 445–475. 10.1113/jp275105.

18. Haemmerle, C.A., Nogueira, M.I., and Watanabe, I.S. (2015). The neural elements in the lining of the ventricular-subventricular zone: making an old story new by high-resolution scanning electron microscopy. Frontiers in neuroanatomy 9, 134. 10.3389/fnana.2015.00134.

19. Møllgård, K., and Wiklund, L. (1979). Serotoninergic synapses on ependymal and hypendymal cells of the rat subcommissural organ. Journal of neurocytology 8, 445–467. 10.1007/bf01214802.

20. Mikkelsen, J.D., Hay-Schmidt, A., and Larsen, P.J. (1997). Central innervation of the rat ependyma and subcommissural organ with special reference to ascending serotoninergic projections from the raphe nuclei. J Comp Neurol 384, 556–568.

21. Simpson, K.L., Fisher, T.M., Waterhouse, B.D., and Lin, R.C. (1998). Projection patterns from the raphe nuclear complex to the ependymal wall of the ventricular system in the rat. J Comp Neurol 399, 61–72. 10.1002/(sici)1096-9861(19980914)399:1<61::aid-cne5>3.0.co;2-8.

22. Murtazina, A.R., Bondarenko, N.S., Pronina, T.S., Chandran, K.I., Bogdanov, V.V., Dilmukhametova, L.K., and Ugrumov, M.V. (2021). A Comparative Analysis of CSF and the Blood Levels of Monoamines As Neurohormones in Rats during Ontogenesis. Acta Naturae 13, 89–97. 10.32607/actanaturae.11516.

23. Shang, P., Ho, A.M., Tufvesson-Alm, M., Lindberg, D.R., Grant, C.W., Orhan, F., Eren, F., Bhat, M., Engberg, G., Schwieler, L., et al. (2022). Identification of cerebrospinal fluid and serum metabolomic biomarkers in first episode psychosis patients. Transl Psychiatry 12, 229 10.1038/s41398-022-02000-1.

24. Verleysdonk, S., Kistner, S., Pfeiffer-Guglielmi, B., Wellard, J., Lupescu, A., Laske, J., Lang, F., Rapp, M., and Hamprecht, B. (2005). Glycogen metabolism in rat ependymal primary cultures: regulation by serotonin. Brain Res 1060, 89–99. 10.1016/j.brainres.2005.08.045.

25. Blanchoin, L., Boujemaa-Paterski, R., Sykes, C., and Plastino, J. (2014). Actin dynamics, architecture, and mechanics in cell motility. Physiol Rev 94, 235–263. 10.1152/physrev.00018.2013.

26. Romanov, R.A., Zeisel, A., Bakker, J., Girach, F., Hellysaz, A., Tomer, R., Alpár, A., Mulder, J., Clotman, F., Keimpema, E., et al. (2017). Molecular interrogation of hypothalamic organization reveals distinct dopamine neuronal subtypes. Nature neuroscience 20, 176–188. 10.1038/nn.4462.

27. Del Bigio, M.R. (2010). Ependymal cells: biology and pathology. Acta neuropathologica 119, 55–73. 10.1007/s00401-009-0624-y.

28. Verkhratsky, A., Semyanov, A., and Zorec, R. (2020). Physiology of Astroglial Excitability. Function (Oxford, England) 1, zqaa016. 10.1093/function/zqaa016.

29. Moriyama, R., Tsukamura, H., Kinoshita, M., Okazaki, H., Kato, Y., and Maeda, K. (2004). In vitro increase in intracellular calcium concentrations induced by low or high extracellular glucose levels in ependymocytes and serotonergic neurons of the rat lower brainstem. Endocrinology 145, 2507–2515. 10.1210/en.2003-1191.

30. Genzen, J.R., Platel, J.C., Rubio, M.E., and Bordey, A. (2009). Ependymal cells along the lateral ventricle express functional P2X(7) receptors. Purinergic Signal 5, 299–307. 10.1007/s11302-009-9143-5.

31. Marichal, N., Fabbiani, G., Trujillo-Cenóz, O., and Russo, R.E. (2016). Purinergic signalling in a latent stem cell niche of the rat spinal cord. Purinergic Signal 12, 331–341. 10.1007/s11302-016-9507-6.

32. Li, Y.C., Bai, W.Z., and Hashikawa, T. (2007). Regionally varying F-actin network in the apical cytoplasm of ependymocytes. Neurosci Res 57, 522–530. 10.1016/j.neures.2006.12.009.

33. Trujillo-Cenóz, O., Rehermann, M.I., Maciel, C., Falco, M.V., Fabbiani, G., and Russo, R.E. (2021). The ependymal cell cytoskeleton in the normal and injured spinal cord of mice. J Neurosci Res 99, 2592–2609. 10.1002/jnr.24918.

34. Veeraval, L., O’Leary, C.J., and Cooper, H.M. (2020). Adherens Junctions: Guardians of Cortical Development. Front Cell Dev Biol 8, 6. 10.3389/fcell.2020.00006.

35. Suzuki, M., Imao, A., Mogami, G., Chishima, R., Watanabe, T., Yamaguchi, T., Morimoto, N., and Wazawa, T. (2016). Strong Dependence of Hydration State of F-Actin on the Bound Mg(2+)/Ca(2+) Ions. J Phys Chem B 120, 6917–6928. 10.1021/acs.jpcb.6b02584.

36. Strzelecka-Golaszewska, H., Wozniak, A., Hult, T., and Lindberg, U. (1996). Effects of the type of divalent cation, C2+ or Mg2+, bound at the high-affinity site and of the ionic composition of the solution on the structure of F-actin. Biochem J 316 (Pt 3), 713-721. 10.1042/bj3160713.

37. Chen, K., Zhang, S., Shao, Y., Guo, M., Zhang, W., and Li, C. (2021). A unique NLRC4 receptor from echinoderms mediates Vibrio phagocytosis via rearrangement of the cytoskeleton and polymerization of F-actin. PLoS Pathog 17, e1010145. 10.1371/journal.ppat.1010145.

38. Zheng, L., Lindsay, A., McSweeney, K., Aplin, J., Forbes, K., Smith, S., Tunwell, R., and Mackrill, J.J. (2022). Ryanodine receptor calcium release channels in trophoblasts and their role in cell migration. Biochim Biophys Acta Mol Cell Res 1869, 119139. 10.1016/j.bbamcr.2021.119139.

39. Fultz, N.E., Bonmassar, G., Setsompop, K., Stickgold, R.A., Rosen, B.R., Polimeni, J.R., and Lewis, L.D. (2019). Coupled electrophysiological, hemodynamic, and cerebrospinal fluid oscillations in human sleep. Science (New York, N.Y.) 366, 628–631. 10.1126/science.aax5440.

40. Li, X., Chen, B., Zhang, D., Wang, S., Feng, Y., Wu, X., Cui, L., Ji, M., Gong, W., Verkhratsky, A., et al. (2023). A novel murine model of mania. Molecular psychiatry. 10.1038/s41380-023-02037-8.

41. Liang, S., Lu, Y., Li, Z., Li, S., Chen, B., Zhang, M., Chen, B., Ji, M., Gong, W., Xia, M., et al. (2020). Iron Aggravates the Depressive Phenotype of Stressed Mice by Compromising the Glymphatic System. Neuroscience bulletin 36, 1542–1546. 10.1007/s12264-020-00539-x.

42. Xia, M., Li, X., Yang, L., Ren, J., Sun, G., Qi, S., Verkhratsky, A., and Li, B. (2017). The ameliorative effect of fluoxetine on neuroinflammation induced by sleep deprivation. Journal of neurochemistry. 10.1111/jnc.14272.

43. Prothmann, C., Wellard, J., Berger, J., Hamprecht, B., and Verleysdonk, S. (2001). Primary cultures as a model for studying ependymal functions: glycogen metabolism in ependymal cells. Brain Res 920, 74–83. 10.1016/s0006-8993(01)03021-9.

44. Li, B., Zhang, S., Zhang, H., Hertz, L., and Peng, L. (2011). Fluoxetine affects GluK2 editing, glutamate-evoked Ca(2+) influx and extracellular signal-regulated kinase phosphorylation in mouse astrocytes. Journal of psychiatry & neuroscience: JPN 36, 322–338. 10.1503/jpn.100094.

45. Hirao, T., Kim, B.G., Habuchi, H., Kawaguchi, K., Nakahari, T., Marunaka, Y., and Asano, S. (2023). Transforming Growth Factor-β1 and Bone Morphogenetic Protein-2 Inhibit Differentiation into Mature Ependymal Multiciliated Cells. Biological & pharmaceutical bulletin 46, 111–122. 10.1248/bpb.b22-00733.

46. Guan, W., Xia, M., Ji, M., Chen, B., Li, S., Zhang, M., Liang, S., Chen, B., Gong, W., Dong, C., et al. (2021). Iron induces two distinct Ca(2+) signalling cascades in astrocytes. Communications biology 4, 525. 10.1038/s42003-021-02060-x.

